# Machine learning on large-scale proteomics data identifies tissue- and cell type-specific proteins

**DOI:** 10.1101/2022.10.02.510525

**Authors:** Tine Claeys, Maxime Menu, Robbin Bouwmeester, Kris Gevaert, Lennart Martens

## Abstract

Using data from 183 public human data sets from PRIDE, a machine learning model was trained to identify tissue and cell-type specific protein patterns. PRIDE projects were searched with ionbot and tissue/cell type annotation was manually added. Data from physiological samples were used to train a Random Forest model on protein abundances to classify samples into tissues and cell types. Subsequently, a one-vs-all classification and feature importance were used to analyse the most discriminating protein abundances per class. Based on protein abundance alone, the model was able to predict tissues with 98% accuracy, and cell types with 99% accuracy. The F-scores describe a clear view on tissue-specific proteins and tissue-specific protein expression patterns. In-depth feature analysis shows slight confusion between physiologically similar tissues, demonstrating the capacity of the algorithm to detect biologically relevant patterns. These results can in turn inform downstream uses, from identification of the tissue of origin of proteins in complex samples such as liquid biopsies, to studying the proteome of tissue-like samples such as organoids and cell lines.

## Introduction

Ongoing advances in mass spectrometry technology result in the field of proteomics being increasingly dominated by big data, which is moreover publicly disseminated through dedicated repositories such as the PRoteomics IDEntifications (PRIDE) database^1,2^. At the time of writing, PRIDE holds more than 16,000 data sets^1,2^, a compendium that holds enormous potential for reuse, reprocessing and repurposing^3^. Examples of this include the training of machine learning tools that predict the liquid chromatography (LC), collisional cross section (CCS), or fragmentation (MS2) behaviour of peptides^4^. In addition to training such method-centric predictors of analyte behaviour however, these data also support more complex biological analyses that rely explicitly on the large scale and heterogeneity of these data^5–8^, such as the discovery of novel protein associations.

Despite the proven utility of public proteomics data re-use, the overall popularity of such approaches remains limited. One reason for this limited uptake of data re-use is the scarcity of key metadata annotations such as tissue type, cell line, treatment, and disease status, amongst others ^9^. Thus, despite the vast amount of publicly available proteomics data, not all data are (re-)usable for in-depth analysis. This substantial hurdle in exploring the potential of proteomics data contradicts the proven interest from the proteomics community in unravelling a (more) holistic view on, for instance, protein expression, function, interaction profile and structure. ^10^ Indeed, major, long running and purpose-designed initiatives such as the Human Proteome Project ^11^ and the Human Protein Atlas ^12^ demonstrate the extensive efforts the community will go to in an attempt to achieve such goals.

Lately, proteomics experiments of healthy human tissues ^13,14^ in their physiological circumstances have lifted the veil on the proteomic composition of these tissues and cell types. The published data from such studies, when combined, creates a treasure trove of information on the human proteome in normal physiological circumstances and should therefore be considered a scientific milestone in the proteomics community. However, each of these data sets comes with its own limitations. As the authors mention^14^, common issues in proteomics studies affect their reproducibility, including (i) limited proteome coverage due to a large amount of proteoforms and their unpredictable expression patterns, (ii) protein inference issues, and (iii) challenges in achieving comprehensive sampling of the proteome. Importantly, however, these situations are precisely where public proteomics data can shine. Indeed, by combining various studies, an overall reduction of such individual, study-specific limitations and biases can be achieved, providing much more solid ground for the discovery of new biological insights^15^.

Here, we re-analysed raw data from 183 PRIDE data sets containing 15,146 raw files that were categorised into 65 human tissues. All data sets were derived from label-free proteomics experiments, devoid of enrichment strategies, and came with unambiguous sample annotation. After further manual curation and annotation on the tissue and cell type level, these data were reprocessed using the ionbot ^16^ search engine. Using the resulting protein identifications, we built a protein expression atlas, hereinafter referred to as PexAt, that was used to train a tissue classifier. Trained exclusively on data from healthy samples in PexAt, this classifier predicts tissue of origin from a proteome with 98% accuracy, and cell type of origin with 99% accuracy. Additionally, the classifier was used to construct a tissue-specific ranking for each protein, alongside a metric for each protein’s importance in identifying each tissue.

## Materials and methods

### Data pre-processing and ionbot reprocessing

In total, 183 publicly available projects from the PRIDE Archive were selected (supplementary table 1). These were selected to only contain samples from human origin without any pre-enrichment of organelles, modified proteins, or protein complexes. These selected projects consisted of 15,146 raw files in total, which were locally reprocessed using ionbot (version 0.6.2) in an open modification search against a protein database containing 75,141 proteins from both Swiss-Prot and TrEMBL (September 2020) as well as common contaminants with S-carbamidomethylation of cysteine and oxidation of methionine as variable modifications ^16^. The raw files were manually annotated for their corresponding tissue and cell type of origin by either the metadata present in PRIDE, or by manual curation of the publication linked to the project. In addition to tissue and cell type, where available, tissue disease status was added, as well as an indication of *in vitro* cell culture growth. Of the 15,146 raw files, 5,441 contained proteomics data from tissues in presumed normal physiological circumstances. The remaining 9,705 files were from diseased samples, fluid samples, cancerous samples, cell cultures, or from ambiguous origin. Cell culture samples were not considered equal to tissue samples due to potential proteomic differences to the tissue of origin caused by differences on a genomic level ^17 18,19^ Fluid samples such as blood and cerebrospinal fluid were excluded from further experiments due to the sample complexity and their tendency to contain so-called tissue leakage proteins from other organs. Peptide identifications were filtered based on three criteria: (i) first ranked PSMs, (ii) q-value ≤ 0.01, and (iii) peptides linked to only one protein. These criteria filtered 905,758,556 PSMs to 35,467,775 PSMs (3.92%), with the unambiguous assignment of PSMs to proteins removing 69,507,260 PSMs. The complete ionbot output, complemented with the available metadata, was stored in a MySQL database, which contained 262,329 unique peptide sequences, irrespective of possible modifications, mapping to 15,108 proteins.

### Filtering and protein quantification

To ensure reliable identification and quantification of proteins, only proteins that were matched by at least three peptides that were each unique linked to this protein were retained. This step reduced the number of proteins by 21.60% to a total of 11,932 identified and quantifiable proteins. These proteins were then quantified using the Normalized Spectral Abundance Factor ^20^, which normalizes spectral counts (*spC)* for a protein *k* by its sequence length (*L*) as counted by amino acid residues, and which further adds an overall normalization for all proteins *N* found in the corresponding raw file *i*:

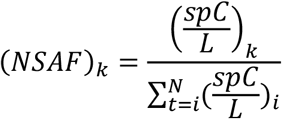

The resulting NSAF expression values were stored in the protein expression atlas (PexAt). Next, a filtered version of this atlas (fPexAt) was obtained by only using proteins present in 90% of the raw files for their corresponding tissue or cell type, similar to a approach used by Kushner et. al ^21^. This reduced the total amount of proteins to 4,169 for the tissue samples and to 4,986 for the cell type samples.

### t-SNE analysis to investigate batch effects

The t-SNE dimensional reduction algorithm was used to investigate possible batch effects between the different experiments due to technical or biological variability. This t-SNE analysis was performed on the physiological fPexAt data. Dimensions were reduced to two components with a perplexity of 15.

### Comparison to the Human Protein Atlas and ProteomicsDB

The NSAF protein expression values were compared to expression values available in the Human Protein Atlas (HPA) and ProteomicsDB ^22^. The HPA data were downloaded from https://www.proteinatlas.org/about/download (number 27, version 21.0) and included the protein expression values (normal_tissue.tsv) and RNA expression values from normal tissues (rna_tissue_hpa.tsv.zip). The former contained ordinal expression values based on immunohistochemistry observations of tissue micro arrays with antibodies, the latter contained transcript expression levels represented by three different metrics: (i) transcripts per million (TPM); (ii) normalized transcripts per million (nTPM); and (iii) protein coding transcripts per million (pTPM). The nTPM metric is commonly used by the HPA to compare gene expression across samples and to classify these genes according to their tissue origin of expression. We here use the same metric for further downstream analyses. ProteomicsDB data (version 4.0) contained the normalized MS1 – iBAQ intensities. Both HPA data and ProteomicsDB data were compared against four variations of the NSAF data: (i) PexAt; (ii) PexAt without the most abundant proteins; (iii) fPexAt; and (iv) fPexAt without the most abundant proteins. The most abundant proteins were here defined as proteins in the top 10% of expression values and were removed to assess their impact on the correlation. An empirical baseline was established by performing the analyses on randomised versions of all four aforementioned NSAF data sets.

The comparison was conducted on two separate levels. First, the correlation on protein level was calculated, based on protein expression values of one protein over various tissues. The PexAt contains 11,879 proteins overlapping with (i) 64.7% of proteins in the HPA antibody data; (ii) 59.9% of proteins in the HPA RNA data; and (iii) 100% of proteins in the PDB data. Second, the correlation on tissue level was calculated, based on protein expression values of various proteins in one tissue. As these tissues show regional differences between sources, e.g. different regions in the brain, they were manually categorized into organs for reliable comparison. The PexAt contains 56 organs overlapping with (i) 86.7% of organs in the HPA antibody data; (ii) 94.9% organs in the HPA RNA data; and (iii) 94.4% organs in the PDB data.

### Training and comparing predictive algorithms

Two training sets were created to train a machine learning based prediction model. These two training sets consisted of either the PexAt or the fPexAt data, but with both only containing data from physiological samples. First, any classes insufficiently represented in the data set were removed. For this, a minimum of three assays was required for each tissue sample. Based on this minimal representation, the following classes were removed: spinal cord, rectum, prostate, placenta, and gall bladder. Applying the same filtering strategy on cell type data led to the removal of the following classes: placenta, spinal cord, skin, prostate, gall bladder and rectum samples. Second, a stratified five-fold cross-validation was repeated five times with different fold. Model performance was determined using accuracy, f1 score, precision and recall.

Five classification algorithms were compared: (i) RandomForest (scikit learn), (ii) Balanced RandomForest (imblearn), (iii) SVM (scikit learn), (iv) XGBClassifier (XGBoost), and (v) LogisiticRegression (scikit learn). As the data contained a varying number of samples per tissue, we tried two methods of class balancing. We compared two methods: (i) set class weight calculated by *weight of a class = (total number of assays – assays of class)/assays of class*, and (ii) balanced mode of the *class_weight* variable using *weight of a class= total number of assays/(number of classes * assays of class)* to the non-balanced performance (supplementary table 2). The highest performing model was selected for further hyperparameter tuning using a GridSearchCV approach.

### OneVsRestClassifier

We used the scikitlearn OneVSRestClassifier with the previously determined highest performing algorithm, the RandomForest algorithm on physiological fPexAt data, to gain class-specific biological insights. Using the same training and testing data, we measured the accuracy of the classifier and used the built-in feature importance module per class for a protein returning the average feature importance over all trees of the RandomForest.

## Results

### Database statistics

The database structure is visualised in **supplementary figure 1** and **supplementary table 3**. The database was built in MySQL and consists of ten tables. The assay table contains the filenames of the 15,146 raw files and has the most connections to other tables. First, a connection with the project table contains general information from PRIDE. Second, the connection with the tissue table contains manually curated information on tissue, cell type, disease status and fluid status. An overview of all tissues and cell types can be found in **supplementary table 4. Supplementary figure 2** shows the number of raw and project files for each tissue. Third, the connection with the peptide table contains the peptide sequence identified by ionbot and links indirectly to the detected protein.

### Atlas statistics

Using the MySQL database, we built two atlases: one that contained all data and one that contained only the filtered data based on 90% protein presence. From each atlas, two sub-atlases were derived: one containing only data collected from healthy tissue samples and one containing all other data (e.g., diseased, cancerous, fluid, cell culture). All four resulting atlases are described in **supplementary table 5**. Although the healthy atlases contain on average only 40% of all raw files, they contain more 1.5 times proteins and 1.8 times more tissues compared to the non-healthy atlases. This is due to the annotation search that mainly focused on healthy tissues and to the availability of some large-scale shotgun proteomics projects on numerous healthy human tissues (PXD010154, PXD020192).

### t-SNE plots

To study batch effects, which are close to inevitable when combining different experiments, a t-SNE analysis on the physiological fPexAt was performed, which visualises the samples in a two-dimensional plot to highlight any clustering between runs. The t-SNE visualisation based on the physiological fPexAt data is shown in **Figure 1** where each sample is coloured according to its cell type. An obvious cell type-dependent clustering is observed and, as expected, some cell types overlap, likely due to similar protein content. For example, pancreas and pancreatic islet cluster together, as do various parts of the eye: retina, sclera, and anterior chamber. However, kidney cell types cluster differently into three defined clusters: (i) podocytes and primary urinary cells, (ii) kidney and single tubules, and (iii) glomeruli.

**Figure 1.**
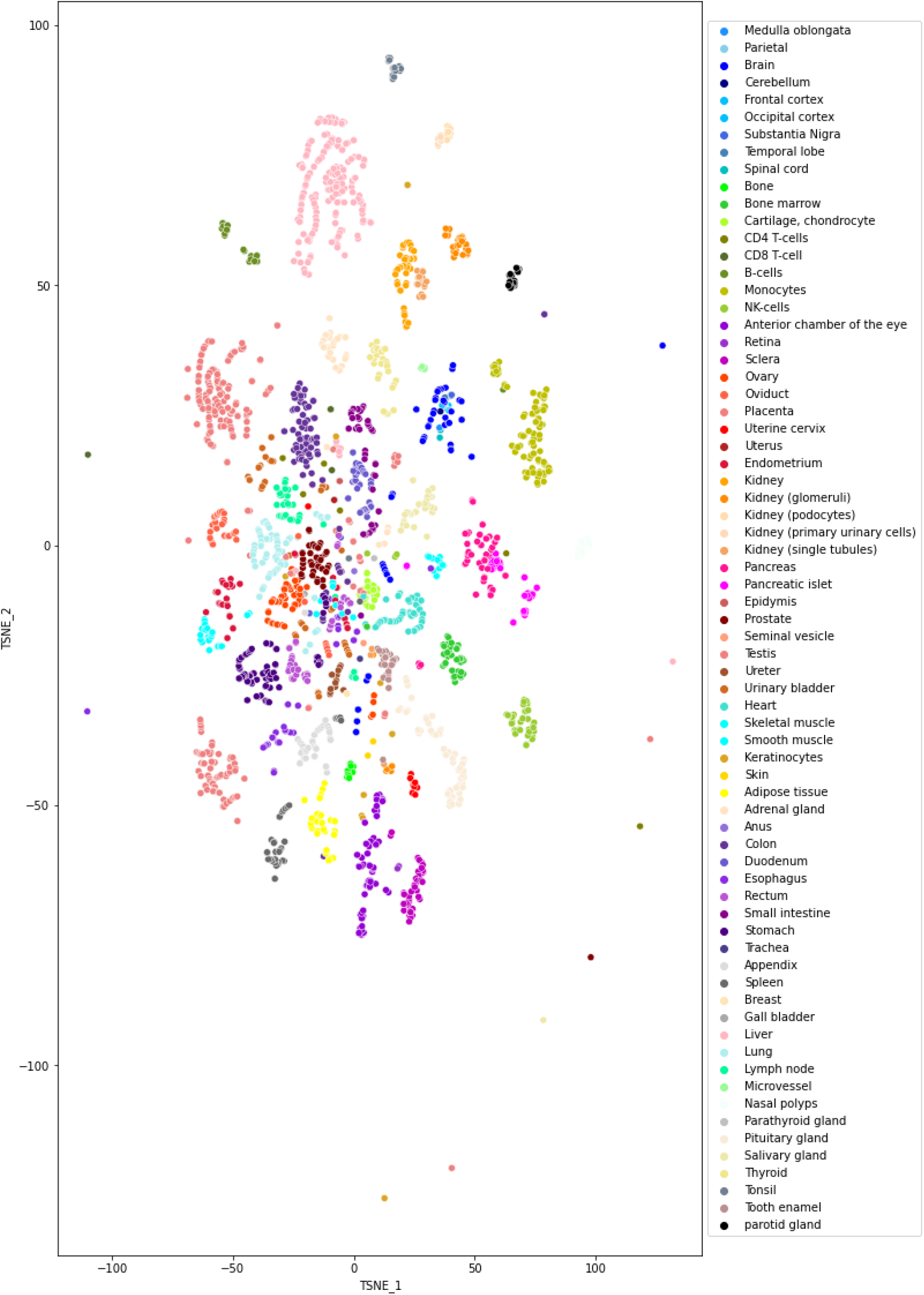
t-SNE visualisation of the physiological fPexAt data. The t-SNE was plotted using two components and a perplexity of 15. Data points were coloured according to cell type.

Cell types that do not cluster despite their similar origin may indicate batch effects. Such effects evoked by outliers were revealed by plotting each individual cell type sample coloured according to its corresponding project. With this approach, outliers in the heart samples were detected. Project 65 (PXD008205) clustered separately from projects 165 (PXD000561), 208 (PXD010154) and 274 (PXD020192). The latter are large-scale projects aimed at proteomic profiling of as many healthy human tissues as possible starting from biopsies, while project 65 used patient-derived induced pluripotent stem cell-derived cardiomyocytes. Hence, the different origin of this sample might explain its outlier position. Another example is seen in the NK cells samples. Here, project 103 (PXD008252) clusters separately from project 165 (PXD000561), possibly due to different NK cell culture conditions in these projects. Although such conditions might affect the NK cell proteome, the samples were retained for analysis to obtain a sufficient amount of training data for the algorithm (**supplementary figure 7**).

The overall picture of this t-SNE analysis shows a strong cell type-dependent clustering. Despite some less dense clusters, our analysis shows that the data are of sufficient quality to proceed with machine learning for further classification refinement.

### Comparison to the Human Protein Atlas and ProteomicsDB

The PexAt data were compared to the Human Protein Atlas and ProteomicsDB, two online sources of protein expression data, with the former relying on in-house generated experimental data, while the latter relies on a subset of public data that is combined at the protein level.

We determined the correlation between the NSAF value and the HPA value by a Kendall rank correlation coefficient. This robust, non-parametric coefficient is particularly useful here as the antibody atlas contains ordinal values, while the protein expression atlas and transcript atlases contain continuous data.

Comparison on the organ level computes the Kendall correlation coefficient per organ. These correlation values were then visualised in a boxplot showing the distribution of all organ-specific Kendall coefficients. The results are visualised in **figure 2** and show the antibody data and nTPM RNA data compared to four variations of the NSAF data and to the randomised versions of these four variations. The first observation is that the median of the antibody atlas fluctuates around 0. This is in contrast to the RNA comparison which has a mean around 0.2 with maxima reaching 0.4. When comparing these values to those of the empirical null values of the randomised data, the ordinal data of the antibody atlas shows a very poor correlation. The RNA data however, which, like the NSAF data, is more fine-grained, shows a strong correlation. The difference in correlation values across atlases is not striking. The fPexAt atlases, containing fewer proteins, show a slightly higher correlation. When computing the protein-specific Kendall correlation coefficients, the boxplots show a more widespread distribution with a larger amount of correlating and anti-correlating values. This is not surprising as there are many more proteins than organs, thus leaving more chances for variation. Similar to the previous comparison, the medians fluctuate around zero, but here the distribution nevertheless seems to be skewed to higher correlation values, especially in the filtered atlases.

**Figure 2.**
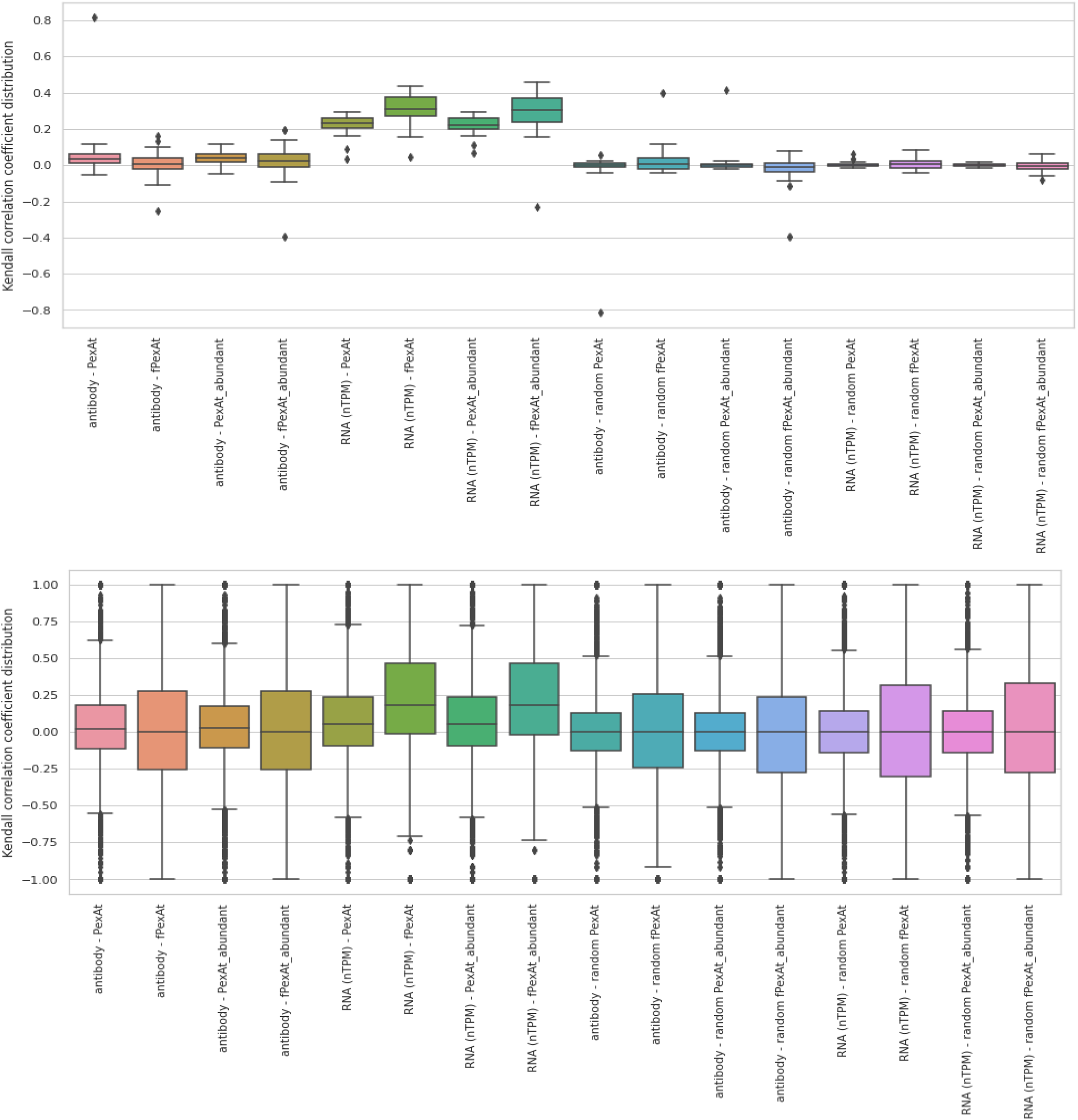
Boxplots visualising the distribution of the Kendall rank correlation coefficient comparison between the Human Protein Atlas data (antibody atlas and normalized RNA atlas (nTPM)) and the true and randomised NSAF atlases (PexAt, fPexAt, PexAt without abundant proteins, fPexAt without abundant proteins) on the level of the organ (top) and the protein (bottom).

We then determined the correlation between the NSAF value and the ProteomicsDB value on organ and protein level by means of the Spearman correlation coefficient. This non-parametric coefficient is particularly useful here because both the NSAF and the ProteomicsDB data contain continuous data but on different scales. The resulting correlation values were then visualised in a boxplot. The correlation on organ level is visualised in **figure 3** and shows that the median varies around 0.6 with minima reaching 0.11 and maxima reaching 0.85. This correlation is very high compared to the randomised data with a median of 0. The fPexAt showed an increased correlation with the ProteomicsDB atlas while removing the abundant proteins had a limited effect. When computing the protein-specific Spearman correlation coefficients, the boxplots show a wide spread, but with a skewed distribution to higher correlation values. The same wide distribution can also be observed in the randomised data sets. However, the random median is centred around 0 while the other medians are centred between 0.20 and 0.30. As the poor protein level correlation can be explained by the variety of metrics for protein expression, and as the organ level correlation shows a positive correlation between the NSAF, HPA and PDB data, we are confident about the quality of our data for machine learning purposes and this mainly because the classifying algorithm will classify samples based on whole proteomes and not individual proteins.

**Figure 3.**
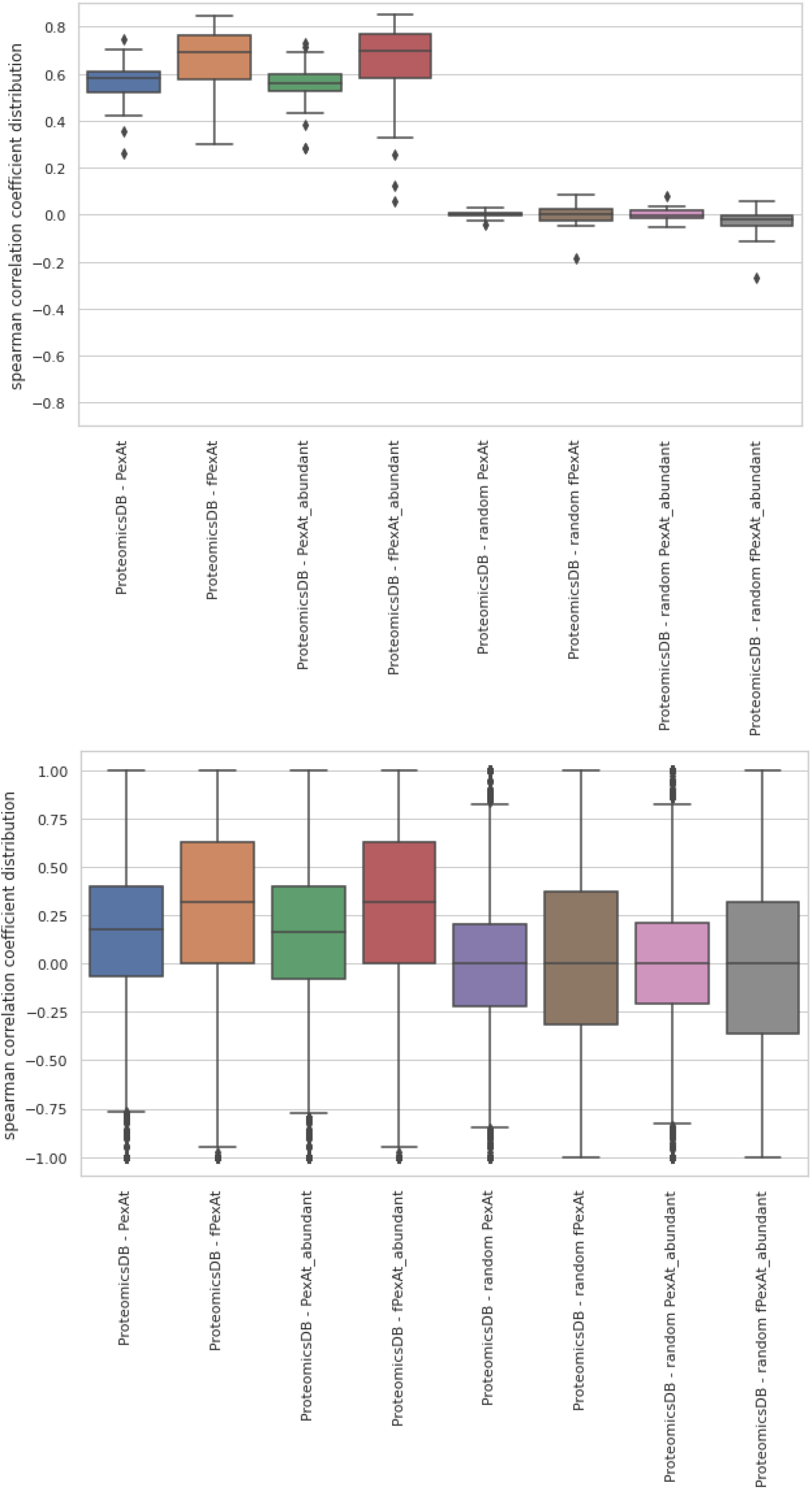
Boxplots visualising the distribution of Spearman correlation coefficient comparison between ProteomicsDB and the true and randomised NSAF atlases (PexAt, fPexAt, PexAt without abundant proteins, fPexAt without abundant proteins) on the level of the organ (top) and the protein (bottom).

### Performance of the algorithms

We compared the performance of 14 different algorithms on four different atlases: (i) tissue PexAt; (ii) tissue fPexAt; (iii) cell-type PexAt, and (iv) cell-type fPexAt. The performance metrics were visualised in a heatmap, resulting in two observations (**supplementary figures 3-6**). First, fPexAt obtained the highest performance. Second, when comparing the different algorithms, the Random Forest algorithm showed the best overall performance. The unbalanced Random Forest algorithm performs the best, but the lack of balancing creates a bias towards prediction of the majority class. Therefore, we opt for the Random Forest algorithm using balanced class weights as the optimal choice and proceeded with optimising the hyperparameters of this predictor using GridSearchCV. The hyperparameter tuned classifiers are the following:

#### Tissue data

Filtered data algorithm:

~~~
RandomForestClassifier(<u>n_estimators=650</u>, max_depth=70, max_features=log2,
bootstrap=False, min_samples_leaf=1, min_samples_split=20, random_state=42,
class_weight=’balanced’)
~~~

Full data algorithm:

~~~
RandomForestClassifier(<u>n_estimators=650</u>, max_depth=90, max_features=’sqrt’,
bootstrap=False, min_samples_leaf=1, min_samples_split=20, random_state=42,
class_weight=’balanced’, n_jobs=-1)
~~~

#### Cell type data

Filtered data algorithm:

~~~
RandomForestClassifier(n_estimators=800, max_depth=70, max_features=’log2’,
bootstrap=False, min_samples_leaf=1, min_samples_split=5, random_state=42,
class_weight=’balanced’, n_jobs=-1)
~~~

Full data algorithm:

~~~
RandomForestClassifier(n_estimators=650, max_depth=50, max_features=’sqrt’,
bootstrap=False, min_samples_leaf=1, min_samples_split=2, random_state=42,
class_weight=’balanced’, n_jobs=-1)
~~~

A comparison of the performance measures, using StratifiedKfold with k=10 gives a thorough overview of the improvement in performance after hyperparameter optimisation. Subsequently, the performance of the final tuned algorithms was visualised using confusion matrices on the test data set (**figures 4** and **5**), which show the improved performance of the fPexAt data set. On the tissue level, while there are some misclassified samples, the large majority is perfectly classified. The misclassifications seem to be related to minority classes such as lymph node being misclassified into majority classes such as eye, thus likely resulting from class imbalance of the data. Misclassification of urinary bladder as esophagus calls for in-depth feature analysis. However, these misclassifications are drastically reduced in the filtered data where the classifier is mostly mistaken by esophagus versus appendix, lymph node versus tonsil, oviduct versus ureter, heart versus cartilage, and ovary versus eye. The latter two can again be explained by the class imbalance, however, all these misclassifications will be further investigated with the feature analysis in the one-vs-rest classifier (see below).

**Figure 4.**
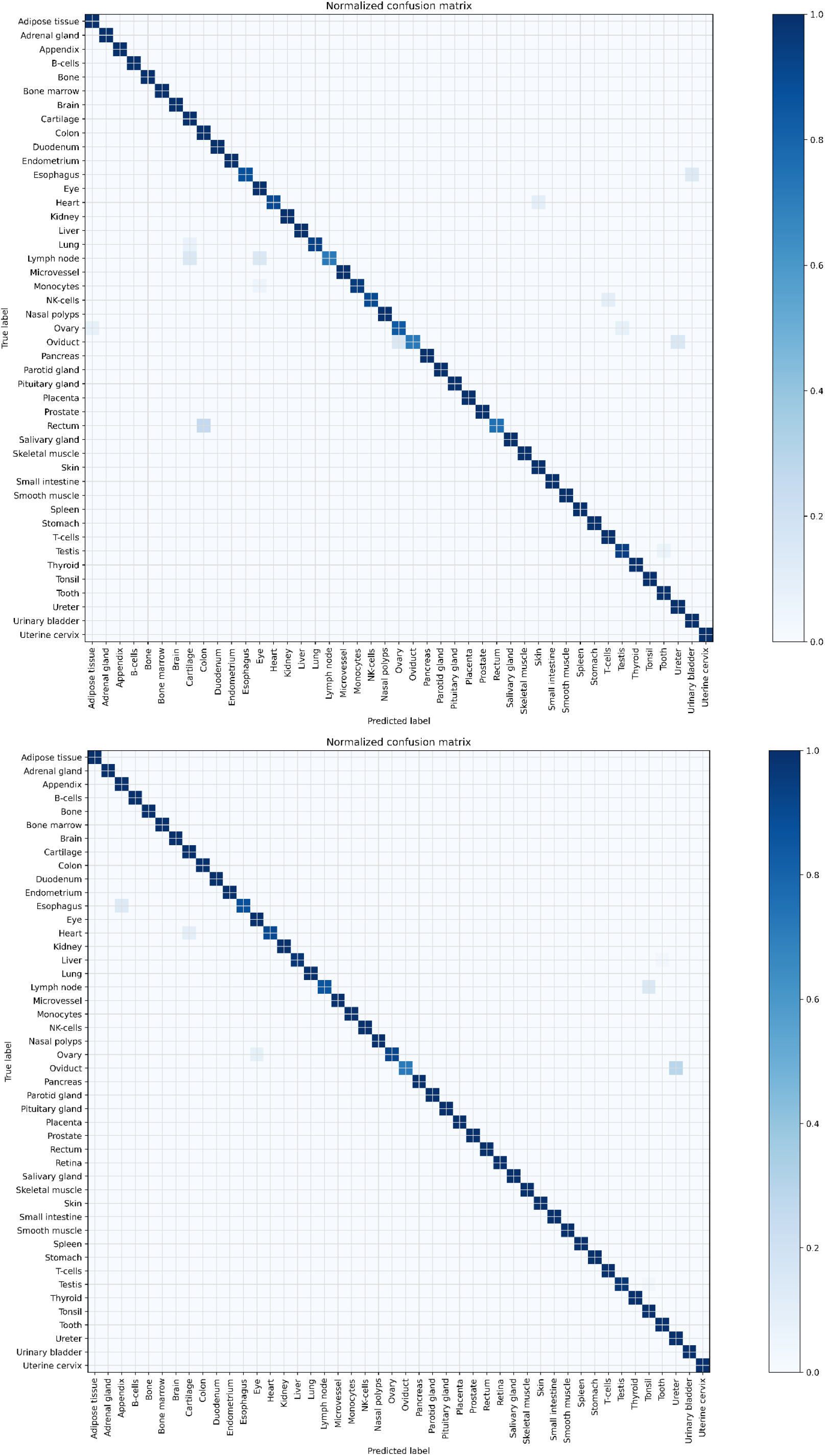
Confusion matrix of the final algorithm on the tissue PexAt (top) and tissue fPexAt (bottom).

**Figure 5.**
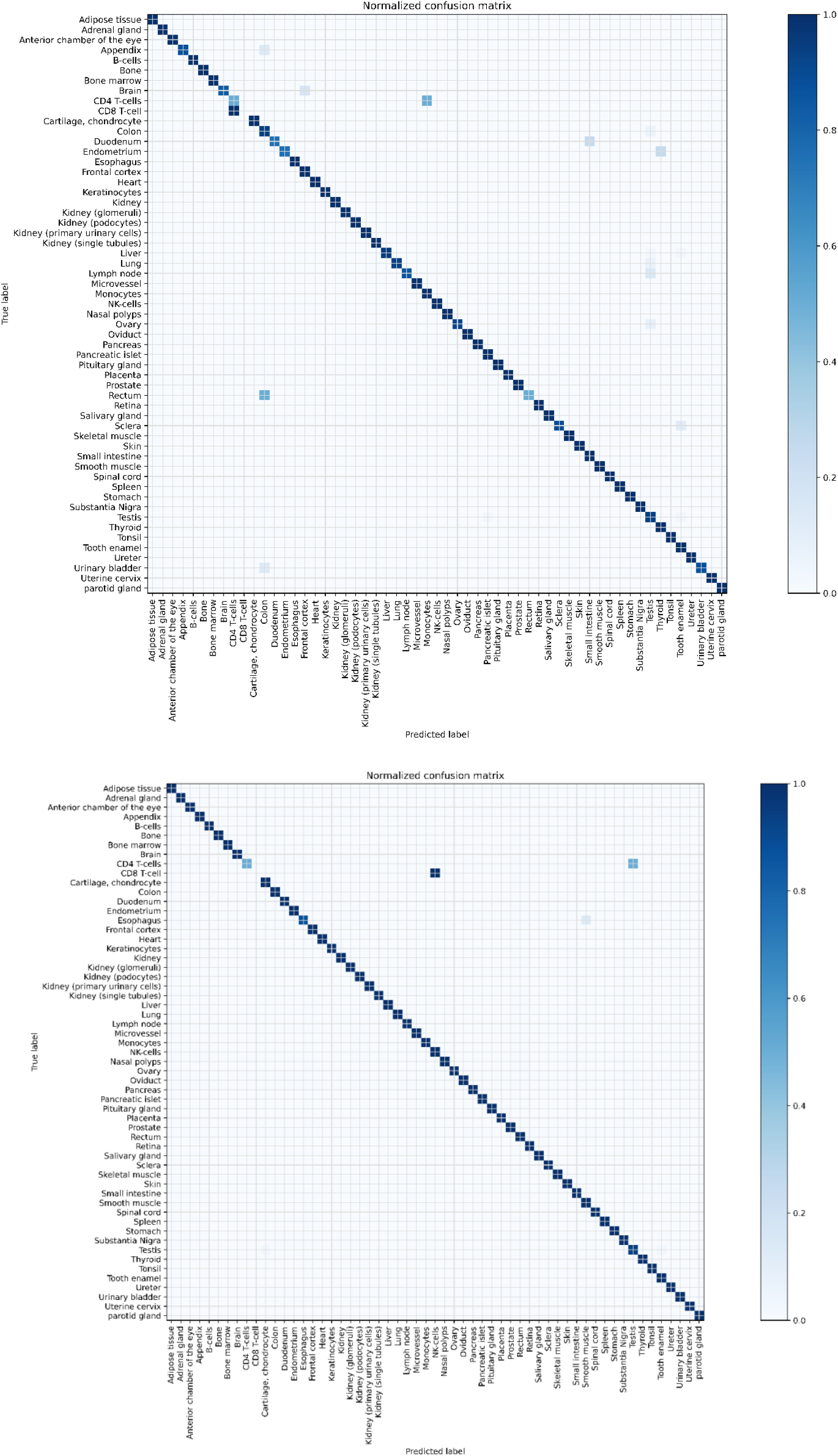
Confusion matrix of the final algorithm on the cell-type PexAt (top) and cell-type fPexAt (bottom).

Also on the cell type level, the fPexAt data set is superior to the complete data set. In the complete data set, several mistakes are made, again on minority classes being predicted as majority classes. Nevertheless, both classifiers are capable of making distinctions between similar cell types such as glomeruli, podocytes and primary urinary cells of the kidney, NK cells and B cells, and also between different brain regions such as the frontal cortex and substantia nigra. A remarkable exception is the inability to correctly classify CD8 T-cells. The algorithm trained on the complete data set classifies these as CD4 T-cells, while the algorithm trained on the filtered data set classifies these as monocytes. The misclassification in itself can likely be explained by the low amount of CD8 data sets.

### Feature analysis of the algorithm

The best performing algorithms, the Random Forest balanced classifier on the fPexAt tissue and cell type data, were used to train a OneVsRestClassifier on the same training and test data. This yielded a 98% classifier accuracy for both the tissue and cell-type data sets. Combining this OneVsRestClassifier with the impurity-based feature importance function of the scikit learn Random Forest determines the importance of a feature to decide if a sample belongs to a specific class. Indeed, this approach provides a metric for judging a protein’s tissue specificity as determined by the classifier. As a RandomForest algorithm combines several decision trees, each protein has a feature importance in all of these trees. Therefore, the interpretable feature importance is a mean of all these values and a standard deviation is added for context clarification. The protein importance thus shows the weight of a protein in the decision of the classifier for the class compared to all other classes (one-vs-rest).

In **supplementary tables 6-9** comparisons are presented of the ten most important proteins between brain classified on the tissue level, and brain, substantia nigra, and frontal cortex all classified on the cell-type level. Enrichment in brain tissue was cross-checked using available public sources such as UniProtKB (literature results), the HPA (protein data) and BGee (gene expression data). The brain classification in both the tissue and cell-type classifiers shows brain-specific proteins with overlap between both classifiers. Moving to cell-type classification, e.g. frontal cortex and substantia nigra, less brain-specific proteins are identified, possibly due to a lower number of samples for these specific brain regions, compared to the larger brain class. Addition of the mean abundance of each protein found in the expression atlas adds information if the presence or absence of the protein plays a role in the tissue classification. For example, in table 5 it becomes clear that the absence of thymosin beta-10 plays a role in the brain classification.

When studying the features for misclassified tissues, some patterns can be observed (**figure 6**). When a misclassification is mainly caused by class imbalance, e.g. ovary *versus* eye, these tissues share a high number of proteins. These shared proteins show a large distribution of feature importances which is contrary to the unique proteins. The proteins that are not shared and therefore unique for their respective tissue all have very low feature importance. This observation validates the assumption that this misclassification was mainly caused by class imbalance. However, in the case of the tonsil *versus* lymphoid node misclassification, the overlap in features is smaller, yet both unique and shared proteins show a wide range of feature importances. In contrast to the previous misclassification, the algorithm is still able to identify relevant tissue-specific proteins despite the confusion. This might be due to the similar nature of lymph nodes and tonsils. Both tissues are secondary lymphoid organs involved in the immune system and have similar functions ^23^, hinting that the overlap in important proteins can therefore be explained by the biological similarity between these two tissues. This shows the biological potential contained in this algorithm and associated feature analyses.

**Figure 6.**
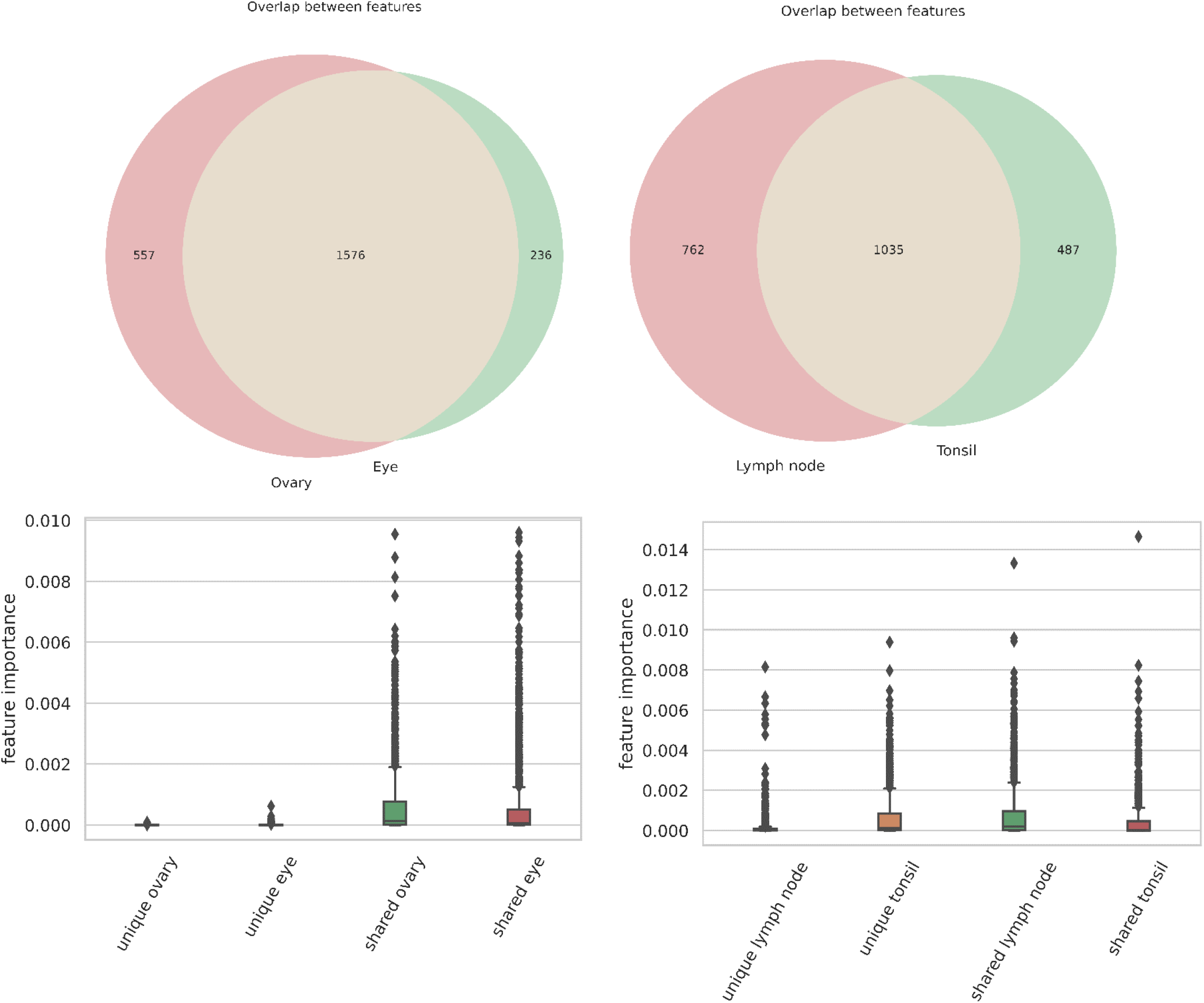
Visualisation of features overlapping in misclassified tissues. Top: absolute number of overlaps between ovary versus eye (left), and lymph node versus tonsil (right). Bottom: feature importances of unique and shared proteins.

## Discussion

Our study combines and reprocesses 183 human proteomics projects from the PRIDE Archive to train a classifier that predicts tissue of origin from a sample based on its proteomic composition with 98% accuracy. The first major bottleneck was gathering the necessary metadata from these experiments. Indeed, the lack of metadata annotation limits the accessibility of large amounts of MS-based proteomics data ^9^. A large manual effort was therefore necessary to assure clean and accurate metadata. Luckily, efforts are ongoing by the HUPO Proteomics Standards Initiative and the ProteomeXchange resource to use the Sample and Data Relationship File (SDRF) for improved metadata annotation of public proteomics data sets in PRIDE ^24^. This important bottleneck will thus hopefully be removed in the future, expanding the opportunities for reprocessing public data.

Reprocessing the raw data with ionbot ^16^ using uniform search settings enabled straightforward re-analysis and comparison. However, this approach comes with two main pitfalls. First, due to the rather generic search settings, experimental differences in set-up or sample preparation are not necessarily optimally represented. Second, as each file is searched separately, the combination of all these experiments would result in the accumulation of a larger than expected number of false positive identifications. This was handled in two ways: (i) only taking proteotypic peptides into account, and (ii) filtering the protein expression atlas by only using proteins present in 90% of the raw files for their corresponding tissue or cell type, hence only taking recurring proteins into account. Despite these precautions, combining various experiments with different experimental setups in different laboratories comes with inevitable batch effects. To observe such effects and, if necessary, adjust data analysis, we used a t-SNE clustering that showed that, in general, similar cell types cluster together. No data adjustment was thus judged necessary.

After constructing the protein expression atlas PexAt, we compared the NSAF expression values to expression values in the HPA and ProteomicsDB databases. This revealed good correlation metrics when comparing protein expression over organs and a widely spread correlation when comparing protein-specific expression. When comparing these to correlation metrics for randomised controls of the NSAF expression values, the same widely spread protein correlation can be observed. Several factors might impact and skew this comparison. First, the overlap in proteins and tissues between both sources is not perfect. Second, tissue level comparison was impacted by the organ classification of tissues, to ensure comparability between the sources. Third, the NSAF values, derived from mass spectrometry experiments, are more fine-grained than the ordinal values in the HPA antibody atlas. Nevertheless, the organ-level data show a strong correlation between the different sources, thereby validating our organ-level use of these data.

Using a Random Forest algorithm, we were able to train a classifier that was able to predict tissue and cell type of a sample with 98% accuracy. Moreover, OneVsRest classification revealed tissue-specific and tissue-important proteins, complete with a metric for their importance in the classification. This result thus shows the biological potential in combining large amounts of public data with machine learning.

The application potential of our study is likely wide-ranging. Drafting tissue proteomes will be an essential component for biomarker identification in liquid biopsies as such studies often search for tissue-leakage proteins. Because we assigned proteins with a metric for their tissue importance, this helps determine the tissue of origin of such tissue-leakage proteins. Additionally, applying the classifier to proteomes of tissue-like samples such as organoids, and immortalised or induced cell lines can be used to reveal their developmental and functional similarity to their tissue of origin. Moreover, this might provide an alternative or complementary approach to the existing methods of tissue and cell line identification such as short tandem repeat profiling ^25^. This also applies for cancerous samples, especially metastases. If a cancerous biopsy reveals a proteomic composition different from the healthy tissue at the biopsy location, this might constitute a clinical marker for metastasis.

Overall, we have shown that proteome-based tissue and cell-type classification is feasible, and that this holds promise for downstream use in typical application areas of proteomics data. Moreover, due to the ever-growing availability of ever more diverse public proteomics data, and due to expected improvements in metadata annotation, the value of such a predictor will only increase over time.

## Acknowledgments

T.C. received funding from the Research Foundation Flanders (FWO) [1S57123N]. R.B. acknowledges funding from the Vlaams Agentschap Innoveren en Ondernemen under project number HBC.2020.2205. K.G. and L. M. acknowledge funding from the European Union’s Horizon 2020 Programme (H2020-INFRAIA-2018-1) [823839]. L. M. acknowledges funding from the Research Foundation Flanders (FWO) [G028821N] and from Ghent University Concerted Research Action [BOF21/GOA/033].

## Data availability

The code used in this work is publicly available at https://github.com/compomics/Tissue_prediction_manuscript. The database and datasets are available via Zenodo at https://doi.org/10.5281/zenodo.7135199. All data are licensed with CC-BY-NC-4.0.

## Supplementary data

**Supplementary table 1.**
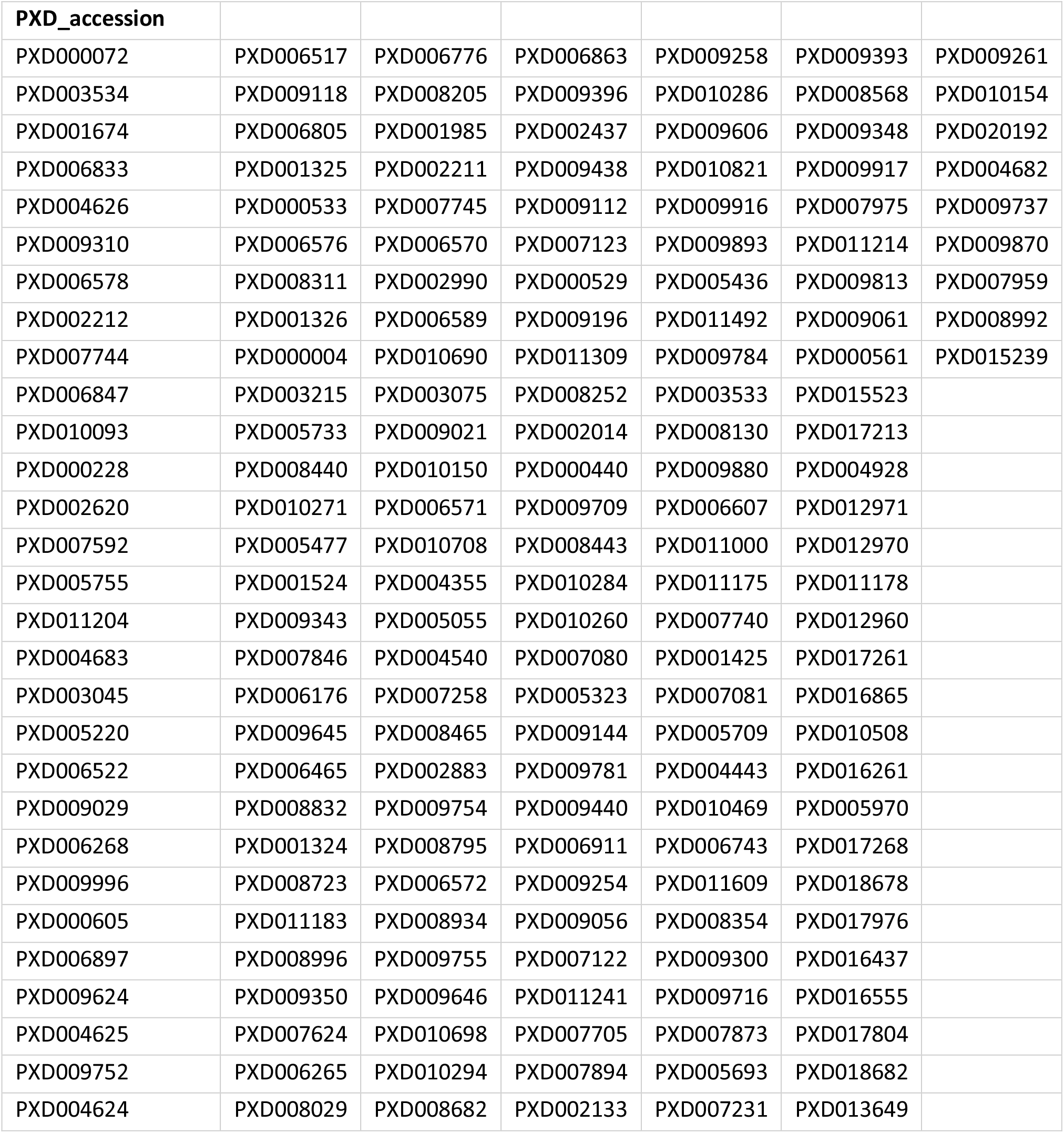
Overview of all used public proteomics datasets and their PRIDE accession number.

| PXD_accession |  |  |  |  |  |  |
| --- | --- | --- | --- | --- | --- | --- |
| PXD000072 | PXD006517 | PXD006776 | PXD006863 | PXD009258 | PXD009393 | PXD009261 |
| PXD003534 | PXD009118 | PXD008205 | PXD009396 | PXD010286 | PXD008568 | PXD010154 |
| PXD001674 | PXD006805 | PXD001985 | PXD002437 | PXD009606 | PXD009348 | PXD020192 |
| PXD006833 | PXD001325 | PXD002211 | PXD009438 | PXD010821 | PXD009917 | PXD004682 |
| PXD004626 | PXD000533 | PXD007745 | PXD009112 | PXD009916 | PXD007975 | PXD009737 |
| PXD009310 | PXD006576 | PXD006570 | PXD007123 | PXD009893 | PXD011214 | PXD009870 |
| PXD006578 | PXD008311 | PXD002990 | PXD000529 | PXD005436 | PXD009813 | PXD007959 |
| PXD002212 | PXD001326 | PXD006589 | PXD009196 | PXD011492 | PXD009061 | PXD008992 |
| PXD007744 | PXD000004 | PXD010690 | PXD011309 | PXD009784 | PXD000561 | PXD015239 |
| PXD006847 | PXD003215 | PXD003075 | PXD008252 | PXD003533 | PXD015523 |  |
| PXD010093 | PXD005733 | PXD009021 | PXD002014 | PXD008130 | PXD017213 |  |
| PXD000228 | PXD008440 | PXD010150 | PXD000440 | PXD009880 | PXD004928 |  |
| PXD002620 | PXD010271 | PXD006571 | PXD009709 | PXD006607 | PXD012971 |  |
| PXD007592 | PXD005477 | PXD010708 | PXD008443 | PXD011000 | PXD012970 |  |
| PXD005755 | PXD001524 | PXD004355 | PXD010284 | PXD011175 | PXD011178 |  |
| PXD011204 | PXD009343 | PXD005055 | PXD010260 | PXD007740 | PXD012960 |  |
| PXD004683 | PXD007846 | PXD004540 | PXD007080 | PXD001425 | PXD017261 |  |
| PXD003045 | PXD006176 | PXD007258 | PXD005323 | PXD007081 | PXD016865 |  |
| PXD005220 | PXD009645 | PXD008465 | PXD009144 | PXD005709 | PXD010508 |  |
| PXD006522 | PXD006465 | PXD002883 | PXD009781 | PXD004443 | PXD016261 |  |
| PXD009029 | PXD008832 | PXD009754 | PXD009440 | PXD010469 | PXD005970 |  |
| PXD006268 | PXD001324 | PXD008795 | PXD006911 | PXD006743 | PXD017268 |  |
| PXD009996 | PXD008723 | PXD006572 | PXD009254 | PXD011609 | PXD018678 |  |
| PXD000605 | PXD011183 | PXD008934 | PXD009056 | PXD008354 | PXD017976 |  |
| PXD006897 | PXD008996 | PXD009755 | PXD007122 | PXD009300 | PXD016437 |  |
| PXD009624 | PXD009350 | PXD009646 | PXD011241 | PXD009716 | PXD016555 |  |
| PXD004625 | PXD007624 | PXD010698 | PXD007705 | PXD007873 | PXD017804 |  |
| PXD009752 | PXD006265 | PXD010294 | PXD007894 | PXD005693 | PXD018682 |  |
| PXD004624 | PXD008029 | PXD008682 | PXD002133 | PXD007231 | PXD013649 |  |

**Supplementary table 2.**
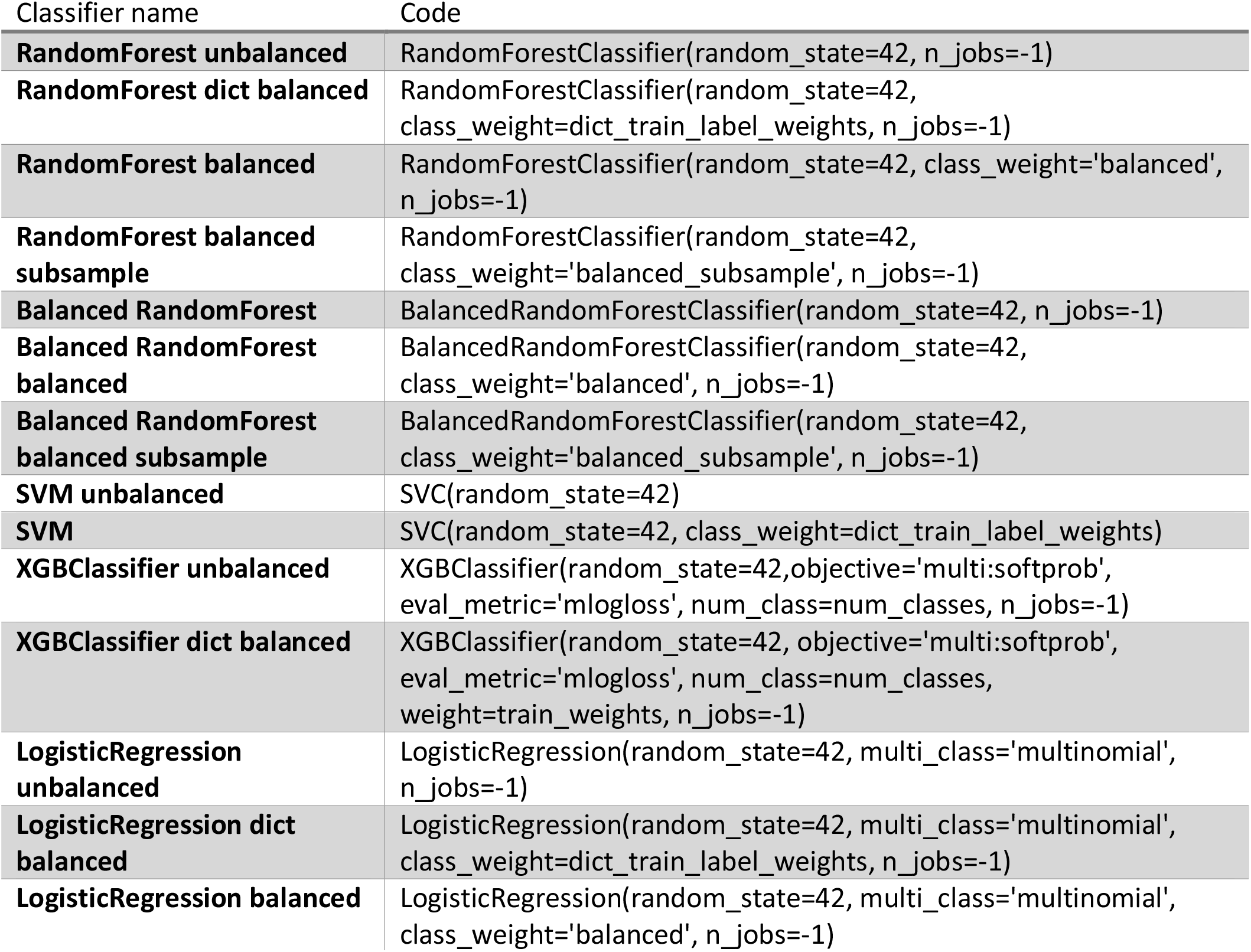
Overview of used algorithms and their hyperparameters.

**Supplementary figure 1.**
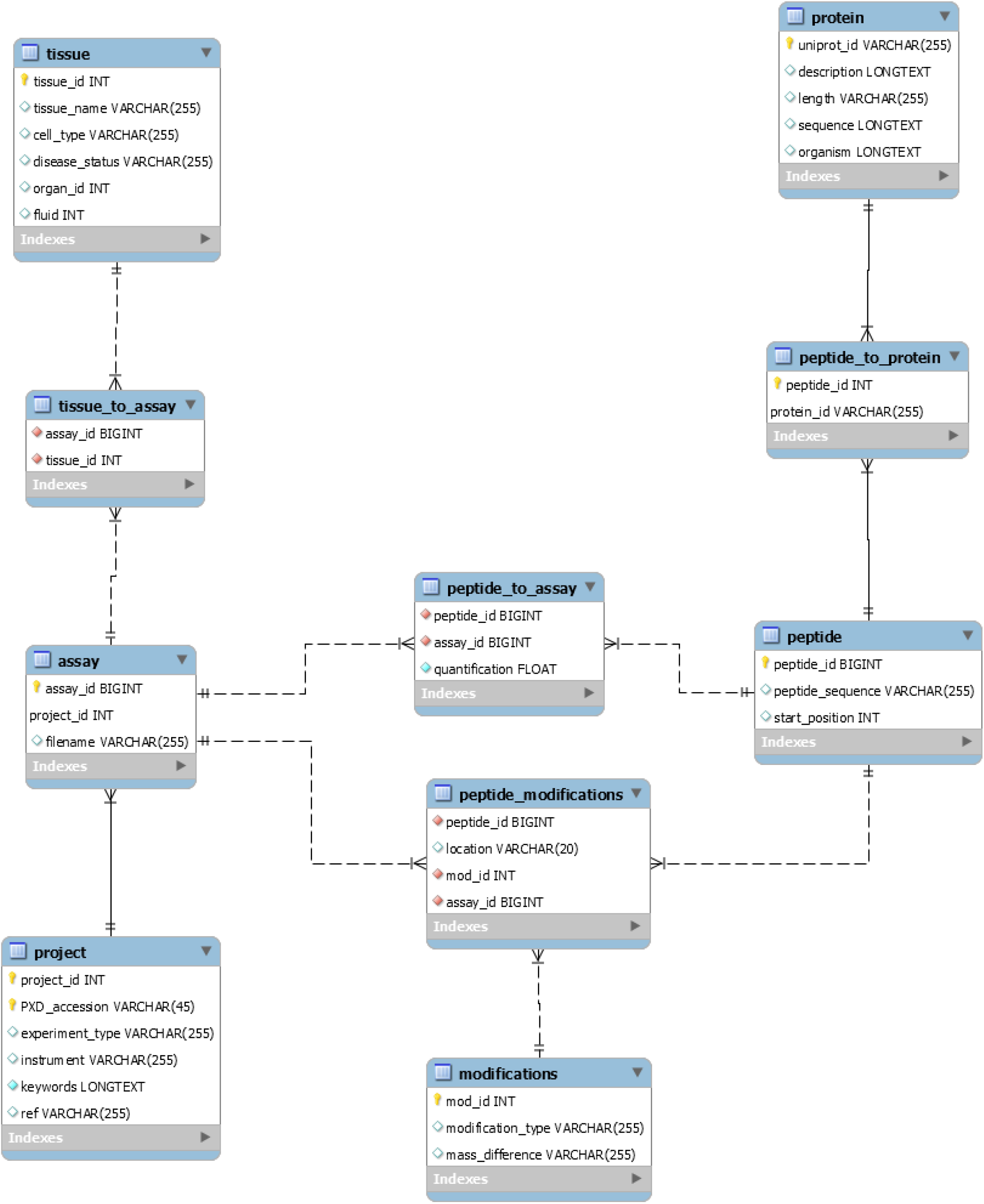
Structure of the protein expression atlas built in MySQL.

**Supplementary table 3.** Structure of the MySQL database with the primary keys and size of each table.

| Table | Primary keys | Rows |
| --- | --- | --- |
| Assay | Assay_id, project_id | 15,146 |
| Modifications | Mod_id | 1,505 |
| Peptide | Peptide_id | 262,329 |
| Peptide_modifications | Peptide_id | 8,020,452 |
| Peptide_to_assay | Assay_id, peptide_id | 14,259,946 |
| Peptide_to_protein | Peptide_id, uniprot_id | 263,466 |
| Project | Project_id, PXD_accession | 183 |
| Protein | Uniprot_id | 15,108 |
| Tissue | Tissue_id | 156 |
| Tissue_to_assay | Assay_id, tissue_id | 15,145 |

**Supplementary table 4.**
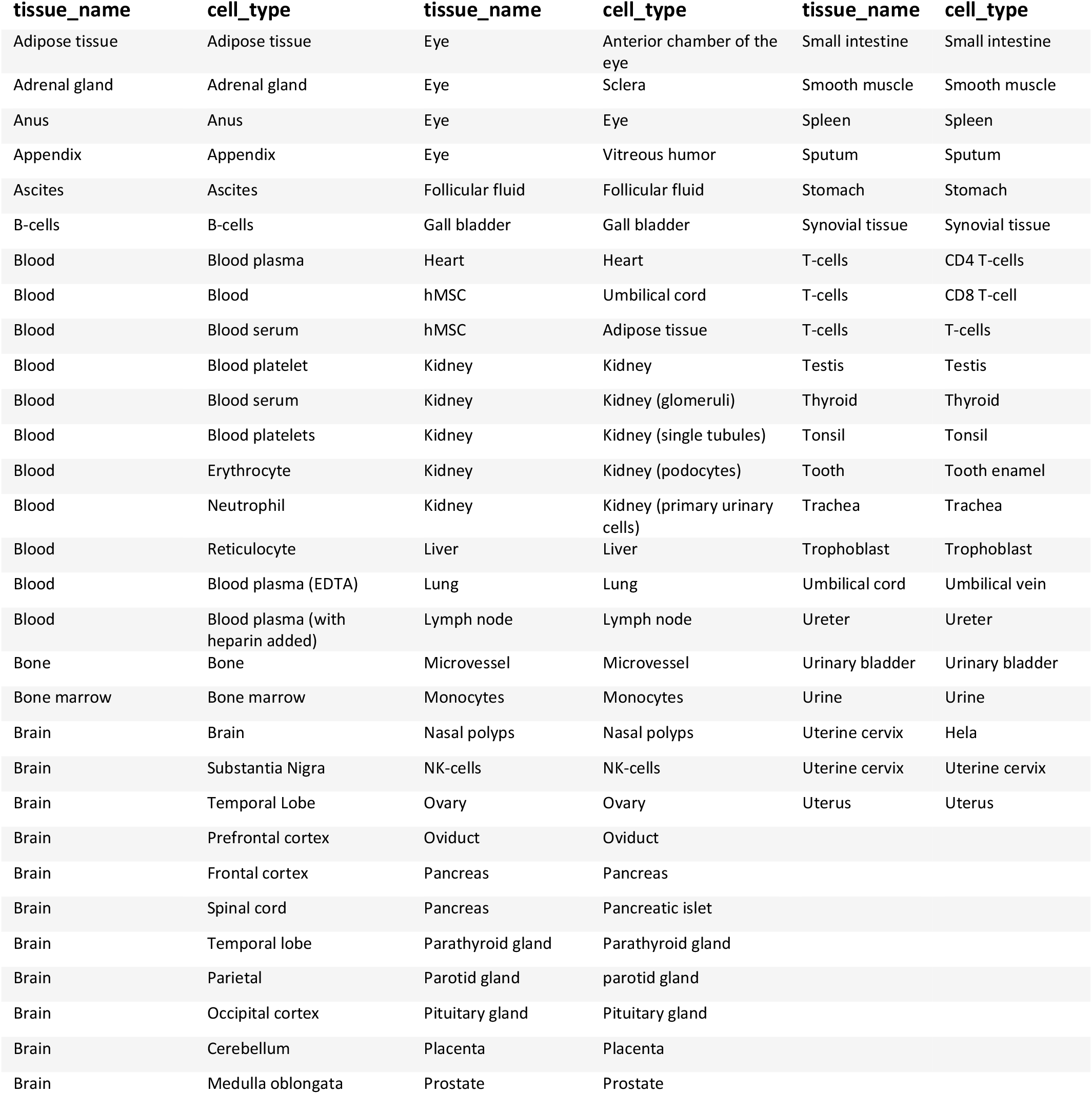
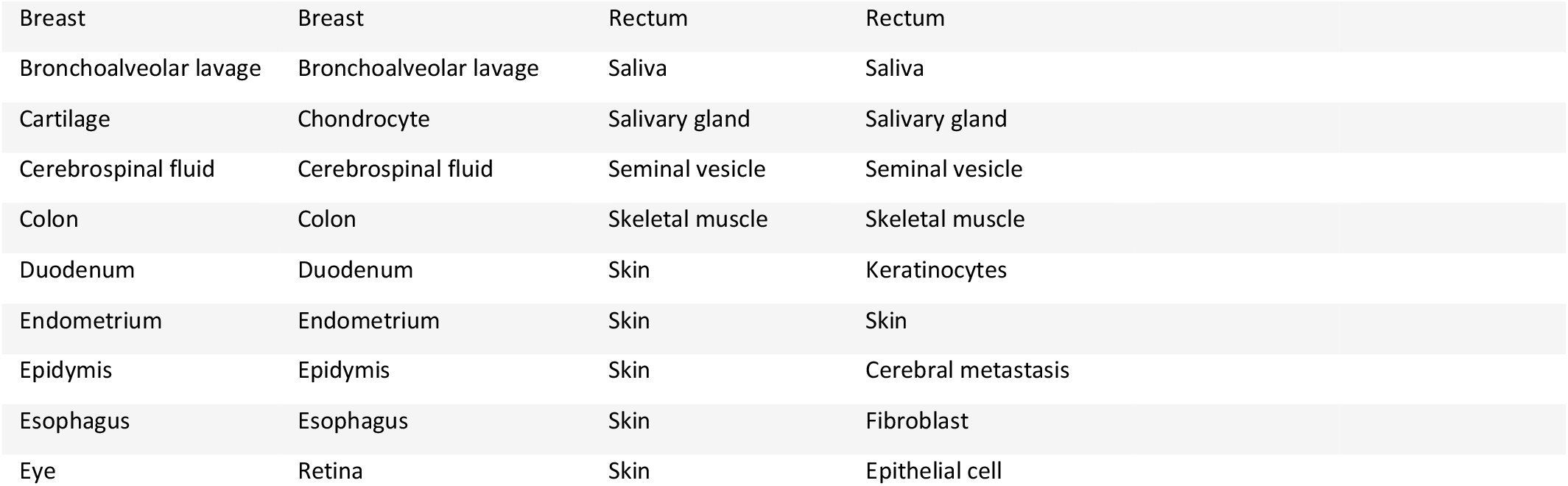
Overview of all tissues and cell types in the database.

| tissue_name | cell_type | tissue_name | cell_type | tissue_name | cell_type |
| --- | --- | --- | --- | --- | --- |
| Adipose tissue | Adipose tissue | Eye | Anterior chamber of the eye | Small intestine | Small intestine |
| Adrenal gland | Adrenal gland | Eye | Sclera | Smooth muscle | Smooth muscle |
| Anus | Anus | Eye | Eye | Spleen | Spleen |
| Appendix | Appendix | Eye | Vitreous humor | Sputum | Sputum |
| Ascites | Ascites | Follicular fluid | Follicular fluid | Stomach | Stomach |
| B-cells | B-cells | Gall bladder | Gall bladder | Synovial tissue | Synovial tissue |
| Blood | Blood plasma | Heart | Heart | T-cells | CD4 T-cells |
| Blood | Blood | hMSC | Umbilical cord | T-cells | CD8 T-cell |
| Blood | Blood serum | hMSC | Adipose tissue | T-cells | T-cells |
| Blood | Blood platelet | Kidney | Kidney | Testis | Testis |
| Blood | Blood serum | Kidney | Kidney (glomeruli) | Thyroid | Thyroid |
| Blood | Blood platelets | Kidney | Kidney (single tubules) | Tonsil | Tonsil |
| Blood | Erythrocyte | Kidney | Kidney (podocytes) | Tooth | Tooth enamel |
| Blood | Neutrophil | Kidney | Kidney (primary urinary cells) | Trachea | Trachea |
| Blood | Reticulocyte | Liver | Liver | Trophoblast | Trophoblast |
| Blood | Blood plasma (EDTA) | Lung | Lung | Umbilical cord | Umbilical vein |
| Blood | Blood plasma (with heparin added) | Lymph node | Lymph node | Ureter | Ureter |
| Bone | Bone | Microvessel | Microvessel | Urinary bladder | Urinary bladder |
| Bone marrow | Bone marrow | Monocytes | Monocytes | Urine | Urine |
| Brain | Brain | Nasal polyps | Nasal polyps | Uterine cervix | Hela |
| Brain | Substantia Nigra | NK-cells | NK-cells | Uterine cervix | Uterine cervix |
| Brain | Temporal Lobe | Ovary | Ovary | Uterus | Uterus |
| Brain | Prefrontal cortex | Oviduct | Oviduct |  |  |
| Brain | Frontal cortex | Pancreas | Pancreas |  |  |
| Brain | Spinal cord | Pancreas | Pancreatic islet |  |  |
| Brain | Temporal lobe | Parathyroid gland | Parathyroid gland |  |  |
| Brain | Parietal | Parotid gland | parotid gland |  |  |
| Brain | Occipital cortex | Pituitary gland | Pituitary gland |  |  |
| Brain | Cerebellum | Placenta | Placenta |  |  |
| Brain | Medulla oblongata | Prostate | Prostate |  |  |

|  |  |  |  |
| --- | --- | --- | --- |
| Breast | Breast | Rectum | Rectum |
| Bronchoalveolar lavage | Bronchoalveolar lavage | Saliva | Saliva |
| Cartilage | Chondrocyte | Salivary gland | Salivary gland |
| Cerebrospinal fluid | Cerebrospinal fluid | Seminal vesicle | Seminal vesicle |
| Colon | Colon | Skeletal muscle | Skeletal muscle |
| Duodenum | Duodenum | Skin | Keratinocytes |
| Endometrium | Endometrium | Skin | Skin |
| Epidymis | Epidymis | Skin | Cerebral metastasis |
| Esophagus | Esophagus | Skin | Fibroblast |
| Eye | Retina | Skin | Epithelial cell |

**Supplementary figure 2.**
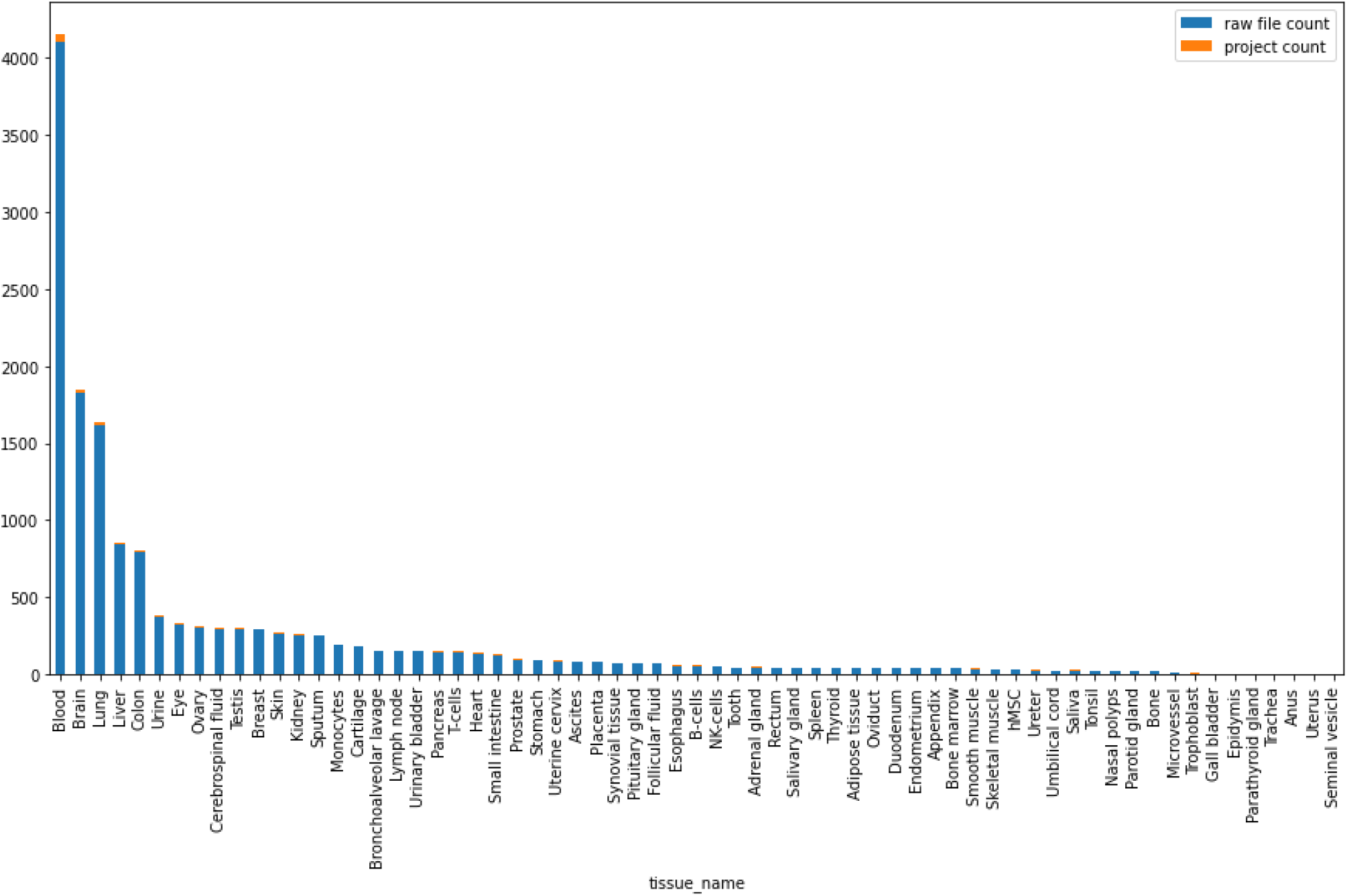
Distribution of the raw files and projects per tissue. The most represented class is blood with 4,105 raw files and the least represented classes are epididymis, parathyroid gland, trachea, anus, uterus and seminal vesicle with only two raw files.

**Supplementary table 5.** Overview of the proteins, raw files, tissues and cell types for the six different atlases that were built using the MySQL database.

| ATLAS | PROTEINS | RAW FILES | TISSUES | CELL TYPES |
| --- | --- | --- | --- | --- |
| COMPLETE ATLAS | 11,891 | 8,279 | 65 | 97 |
| COMPLETE ATLAS HEALTHY TISSUE | 11,809 | 3,191 | 58 | 81 |
| COMPLETE ATLAS DISEASED TISSUE | 11,538 | 5,088 | 32 | 45 |
| FILTERED ATLAS | 5,110 | 8,246 | 65 | 97 |
| FILTERED ATLAS HEALTHY TISSUE | 5,021 | 3,173 | 58 | 81 |
| FILTERED ATLAS DISEASED TISSUE | 3,371 | 5,073 | 32 | 45 |

**Supplementary figure 3.**
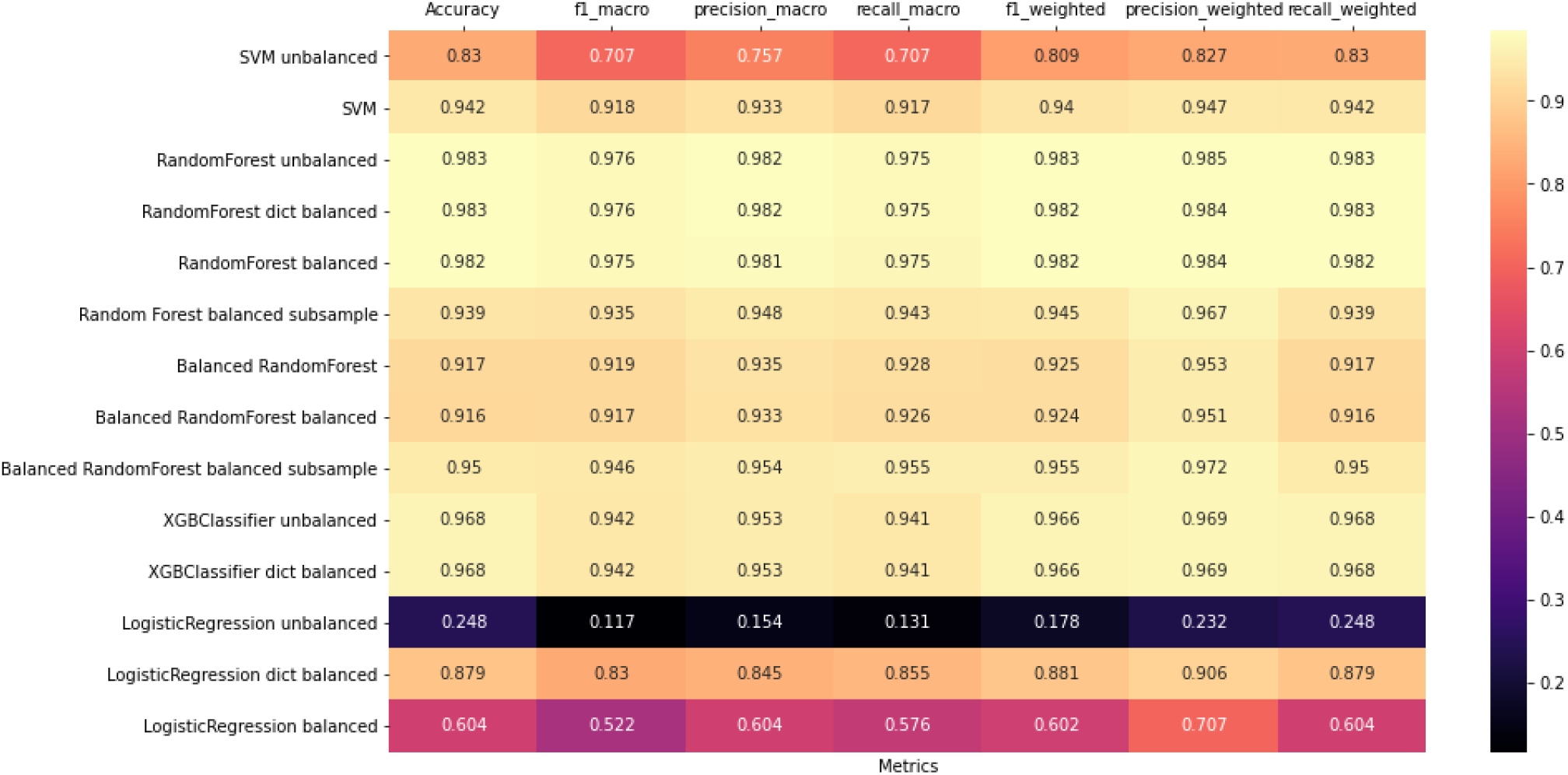
Heatmap visualising classifier performance on tissue fPexAt data

**Supplementary figure 4.**
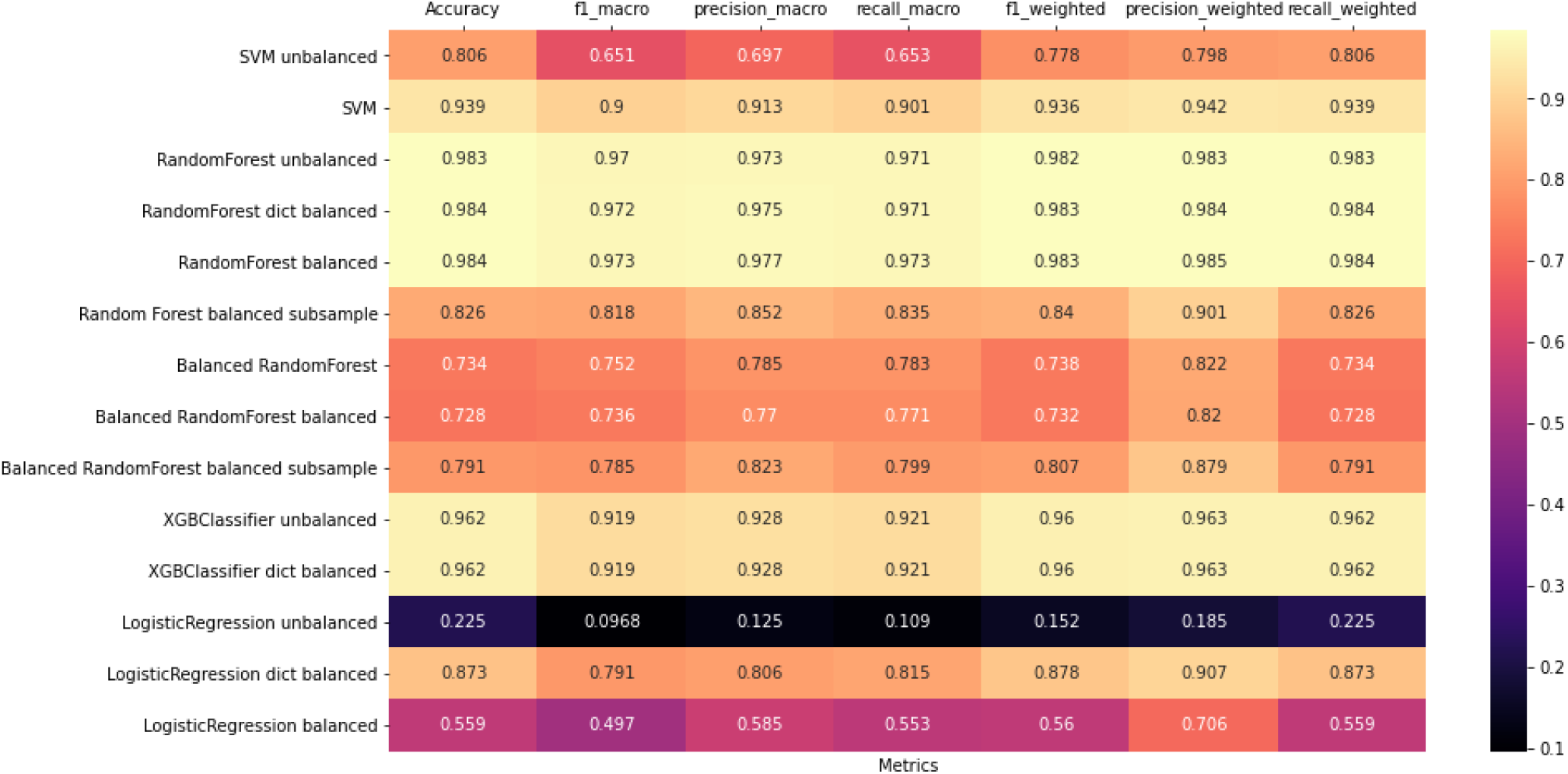
Heatmap visualising classifier performance on cell-type fPexAt data

**Supplementary figure 5.**
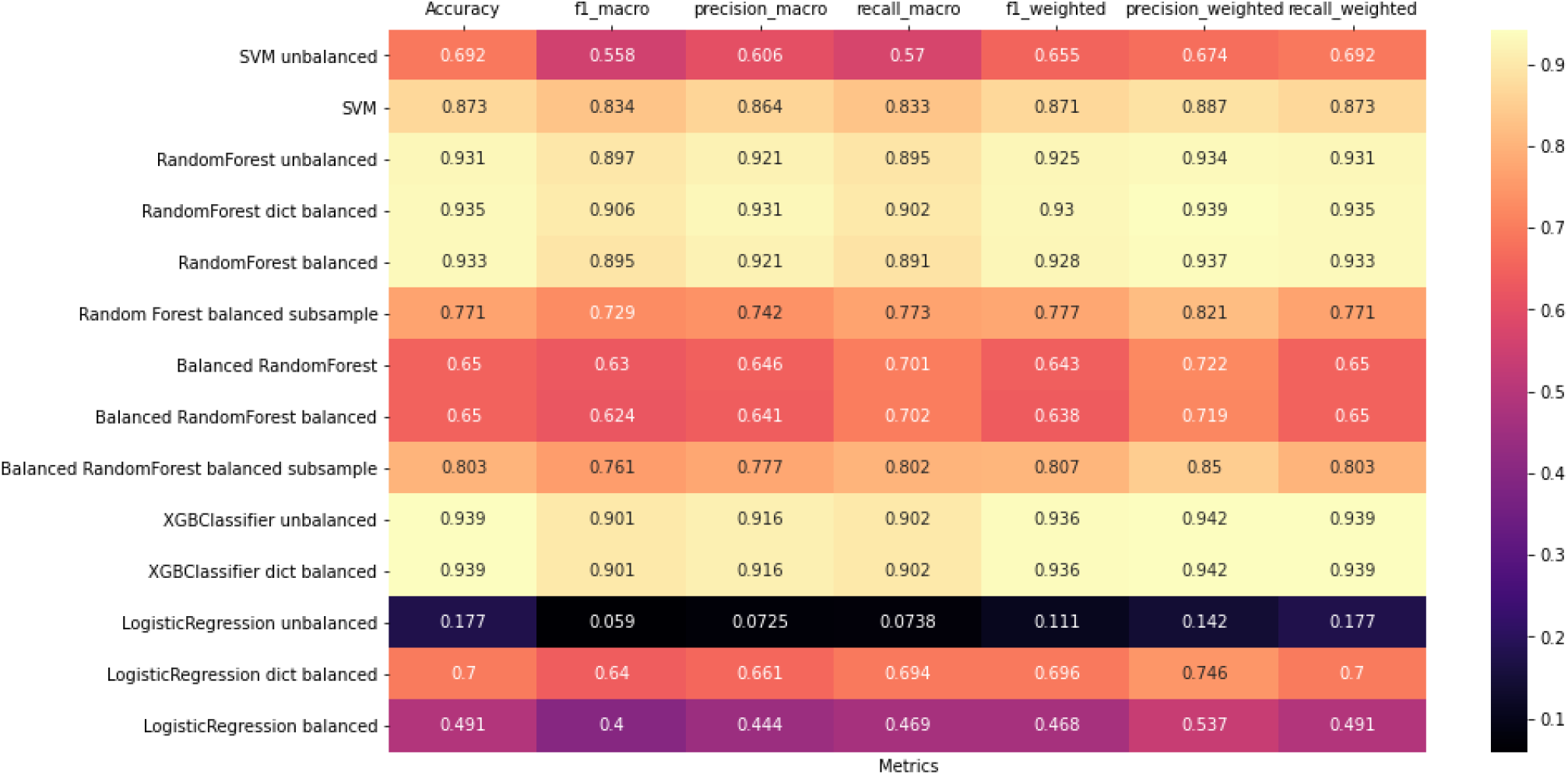
Heatmap visualising classifier performance on the tissue PexAt data.

**Supplementary figure 6.**
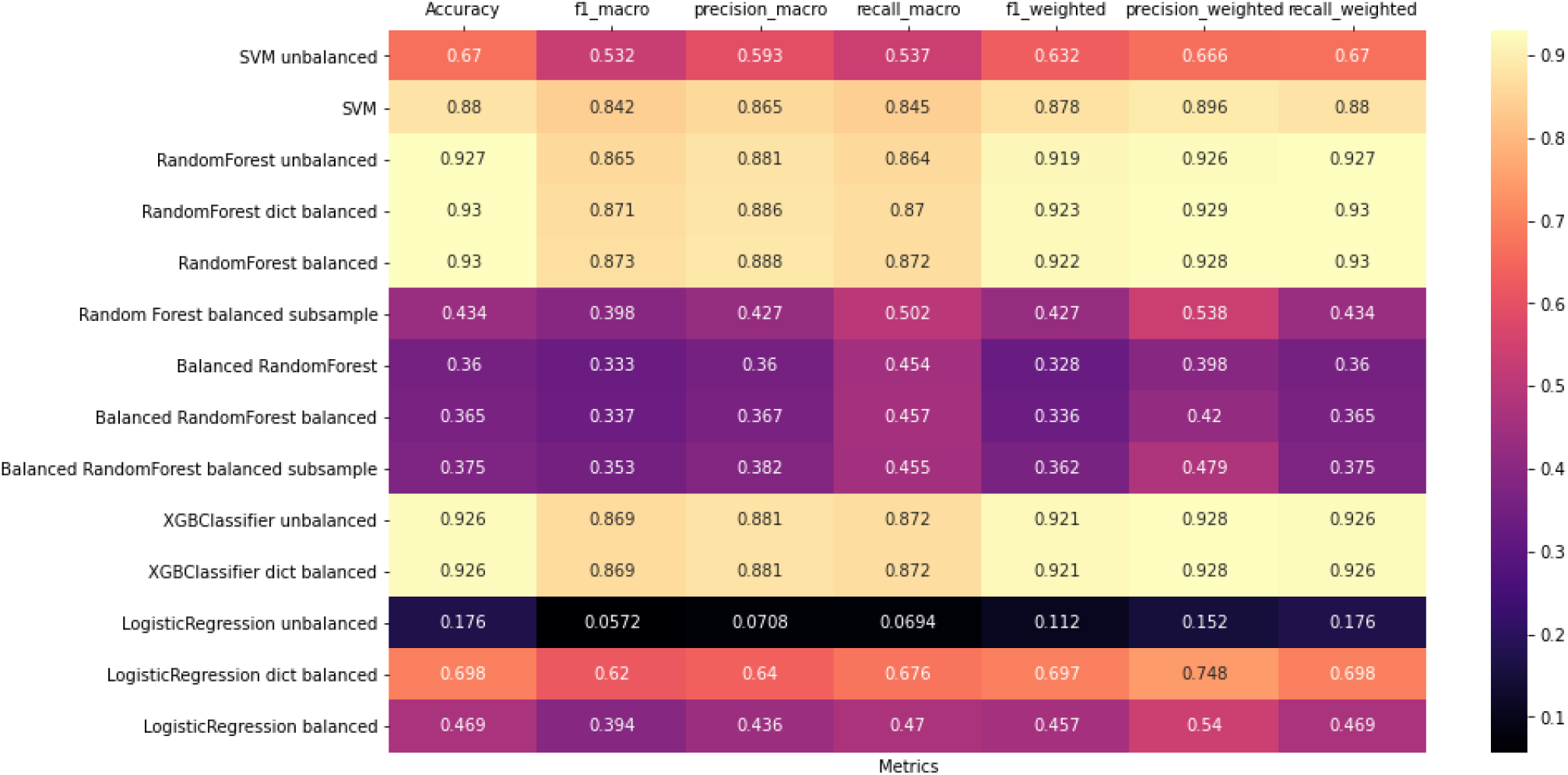
Heatmap visualising classifier performance on the cell-type PexAt data

**Supplementary table 6.** The ten most important proteins for brain classification in the tissue classifier with their mean importance, the standard deviation over all decision trees. The column ‘Brain enrichment’ contains a surface-level cross-check with three other sources: UniProtKB, the Human Protein Atlas (HPA) and the Bgee gene expression database.

Brain - tissue classifier
| uniprot ID | importance | standard deviation | mean abundance | protein name | Brain enrichment |
| --- | --- | --- | --- | --- | --- |
| **Q16352** | 0.0139 | 0.0651 | 0.0161 | Alpha-internexin | yes, all sources |
| **P80723** | 0.0116 | 0.0409 | 0.0218 | Brain acid soluble protein 1 | yes, all sources |
| **P62328** | 0.0093 | 0.0239 | 0.0520 | Thymosin beta-4 | yes, Bgee (Uniprot and HPA: in blood) |
| **Q12860** | 0.0085 | 0.0717 | 0.0094 | Contactin-1 | yes, all sources |
| **Q96GW7** | 0.0082 | 0.0610 | 0.0020 | Brevican core protein | yes, all sources |
| **P63313** | 0.0080 | 0.0212 | 0.0000 | Thymosin beta-10 | yes, Bgee (Uniprot and HPA: in blood) |
| **P07196** | 0.0079 | 0.0577 | 0.0073 | Neurofilament light polypeptide (NF-L) | yes, all sources |
| **P35612** | 0.0072 | 0.0608 | 0.0006 | Beta-adducin | yes, all sources |
| **O94811** | 0.0072 | 0.0601 | 0.0041 | Tubulin polymerization-promoting protein | yes, all sources |
| **Q7L0J3** | 0.0072 | 0.0595 | 0.0026 | Synaptic vesicle glycoprotein 2A | yes, all sources |

**Supplementary table 7.** The ten most important proteins for brain classification in the cell type classifier with their mean importance, the standard deviation over all decision trees. The column ‘Brain enrichment’ contains a surface-level cross-check with three other sources: UniProtKB, the Human Protein Atlas (HPA) and the Bgee gene expression database.

Brain - cell classifier
| uniprot ID | importance | standard deviation | mean abundance | protein name | Brain enrichment |
| --- | --- | --- | --- | --- | --- |
| **Q16352** | 0.037583 | 0.12124 | 0.016075 | Alpha-internexin | yes, all sources |
| **P07196** | 0.026597 | 0.123581 | 0.007261 | Neurofilament light polypeptide | yes, all sources |
| **P78357** | 0.022235 | 0.111044 | 0.00183 | Contactin-associated protein 1 | yes, all sources |
| **Q16720** | 0.018581 | 0.0992 | 0.000913 | Plasma membrane calcium-transporting ATPase 3 | yes, all sources |
| **P80723** | 0.018152 | 0.052642 | 0.021842 | Brain acid soluble protein 1 | yes, all sources |
| **P17600** | 0.017851 | 0.089281 | 0.004646 | Synapsin-1 | yes, all sources |
| **P60201** | 0.016725 | 0.090625 | 0.050548 | Myelin proteolipid protein | yes, all sources |
| **Q12860** | 0.016487 | 0.100304 | 0.009409 | Contactin-1 | yes, all sources |
| **P05129** | 0.016079 | 0.084747 | 0.000902 | Protein kinase C gamma type | yes, all sources |
| **Q92752** | 0.015677 | 0.092039 | 0.008958 | Tenascin-R | yes, all sources |

**Supplementary table 8.**
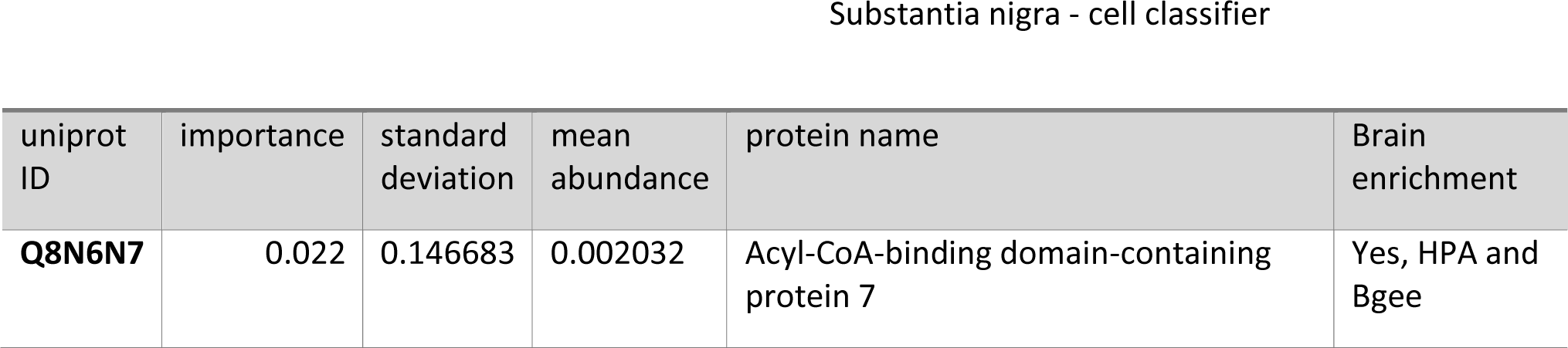
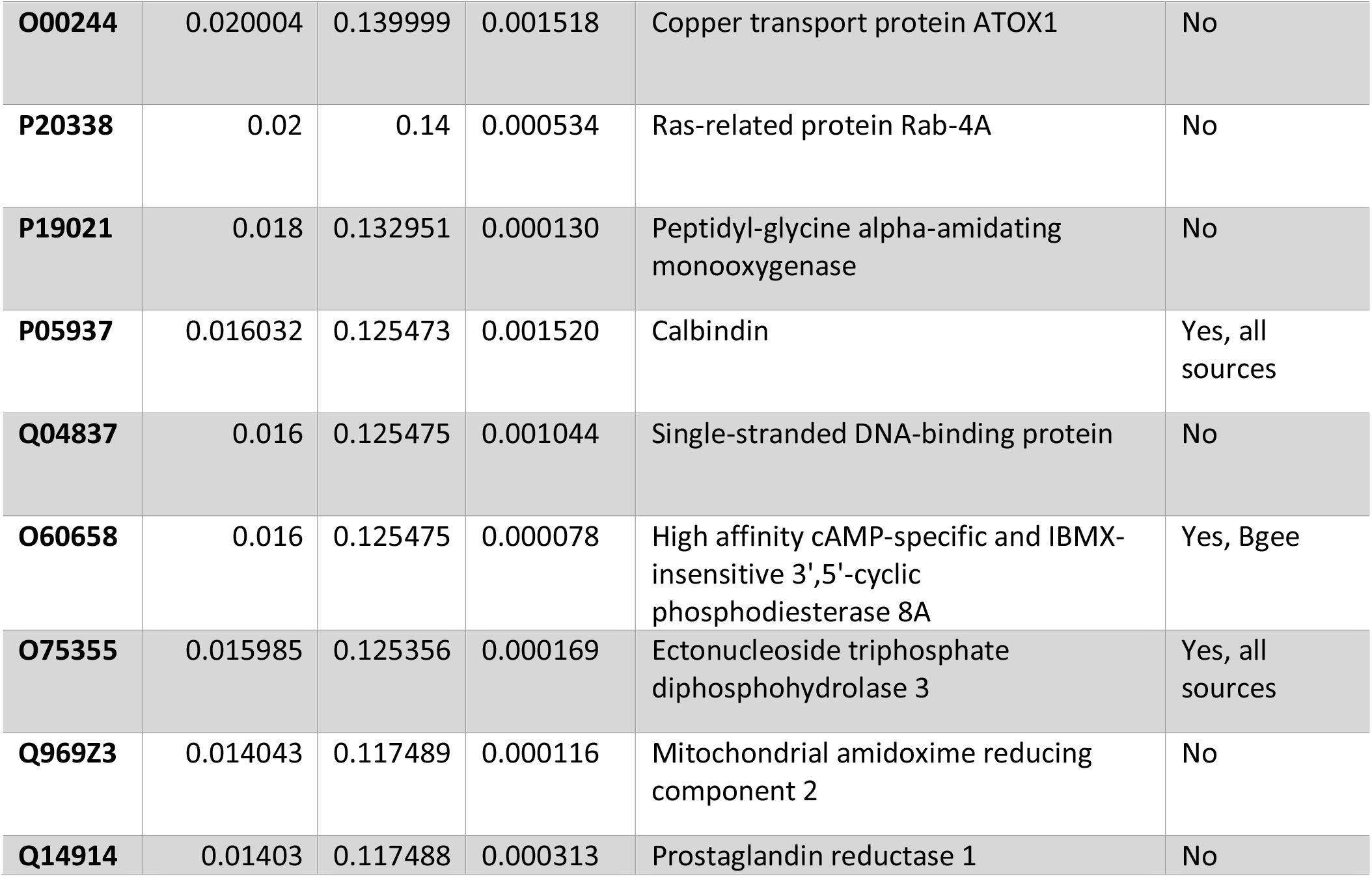
The ten most important proteins for substantia nigra classification in the cell type classifier with their mean importance, the standard deviation over all decision trees. The column ‘Brain enrichment’ contains a surface-level cross-check with three other sources: UniProtKB, the Human Protein Atlas (HPA) and the Bgee gene expression database.

**Supplementary table 9.** The ten most important proteins for frontal cortex classification in the cell type classifier with their mean importance, the standard deviation over all decision trees. The column ‘Brain enrichment’ contains a surface-level cross-check with three other sources: UniProtKB, the Human Protein Atlas (HPA) and the Bgee gene expression database.

| uniprot ID | importance | standard deviation | mean abundance | protein name | Brain enrichment |
| --- | --- | --- | --- | --- | --- |
| **P42684** | 0.030019 | 0.170584419 | 0.0000245 | Tyrosine-protein kinase ABL2 | No |
| **Q9UNA1** | 0.02 | 0.14 | 0.0000522 | Rho GTPase-activating protein 26 | No |
| **Q6ICH7** | 0.018 | 0.132951119 | 0.0000757 | Aspartate beta-hydroxylase domain-containing protein 2 | yes, all sources |
| **Q5TCQ9** | 0.01604 | 0.125472028 | 0.0000176 | Membrane-associated guanylate kinase, WW and PDZ domain-containing protein 3 | No |
| **Q9NVE7** | 0.016 | 0.125475097 | 0.0000359 | 4'-phosphopantetheine phosphatase | No |
| **P10915** | 0.015987 | 0.125355759 | 0.0004557 | Hyaluronan and proteoglycan link protein 1 | Yes |
| **O96007** | 0.014043 | 0.117489241 | 0.0001154 | Molybdopterin synthase catalytic subunit | yes, all sources |
| **Q9NZ56** | 0.014025 | 0.117488168 | 0.0001073 | Formin-2 | yes, all sources |
| **Q8IXS8** | 0.014021 | 0.117488866 | 0.0000485 | Protein FAM126B | Yes, Bgee |
| **Q96AB3** | 0.014 | 0.117490425 | 0.0000407 | Isochorismatase domain-containing protein 2 | No |

**Supplementary figure 7.**
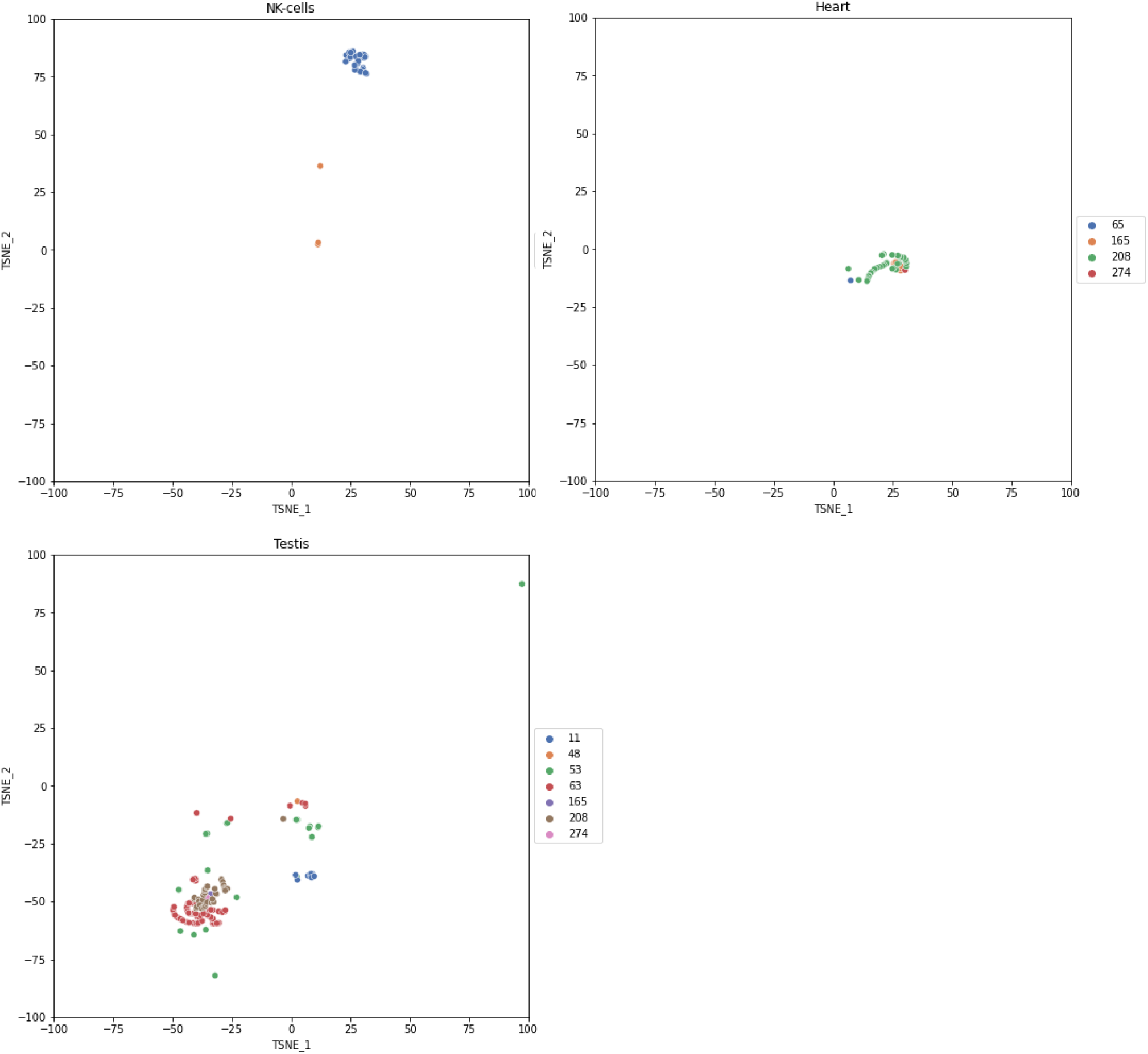
t-SNE visualisation of NK-cells, heart and testis, coloured according to the assays.

**Supplementary figure 8.**
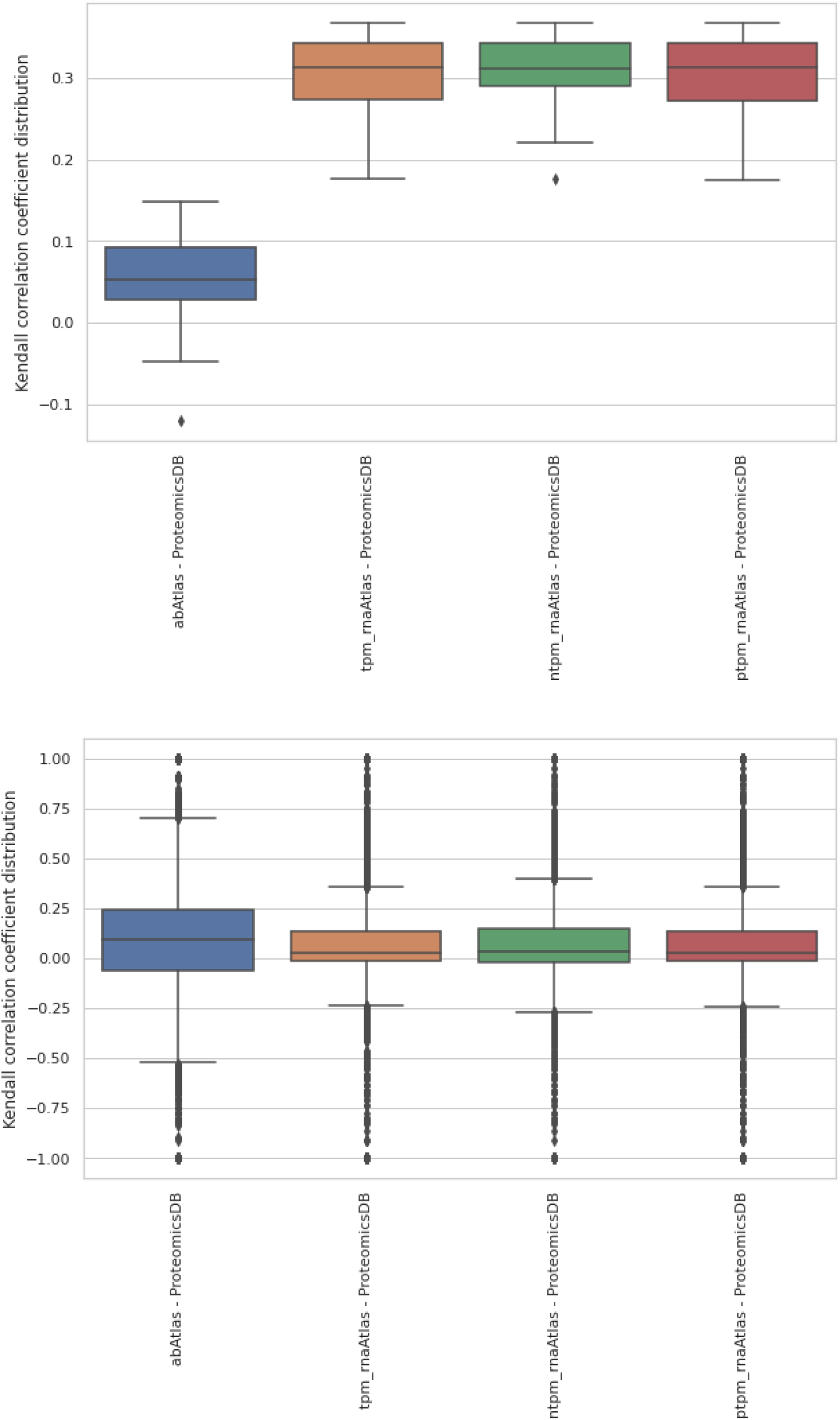
Boxplots visualising the distribution of Kendall correlation coefficient comparison between ProteomicsDB and the Human Protein Atlas data (antibody atlas, unnormalized RNA atlas (tpm), normalized RNA atlas (nTPM) and protein-coding RNA atlas (ptpm) on the level of the organ (top) and the protein (bottom).

